# Structural insights into the modes of relaxin-binding and tethered-agonist activation of RXFP1 and RXFP2

**DOI:** 10.1101/2021.06.06.446989

**Authors:** Ashish Sethi, Shoni Bruell, Tim Ryan, Fei Yan, Mohammad Tanipour, Yee-Foong Mok, Chris Draper-Joyce, Yogesh Khandokar, Riley D. Metcalfe, Michael D. W. Griffin, Daniel J. Scott, Mohammad Akhter Hossain, Emma J. Petrie, Ross A. D. Bathgate, Paul R. Gooley

## Abstract

Our poor understanding of the mechanism by which the peptide-hormone H2 relaxin activates its G protein coupled receptor, RXFP1 and the related receptor RXFP2, has hindered progress in its therapeutic development. Both receptors possess large ectodomains, which bind H2 relaxin, and contain an N-terminal LDLa module that is essential for receptor signalling and postulated to be a tethered agonist. Here, we show that a conserved motif (GDxxGWxxxF), C-terminal to the LDLa, is critical for receptor activity. Importantly, this motif adopts different structures in RXFP1 and RXFP2, suggesting distinct activation mechanisms. For RXFP1, the motif is flexible, weakly associates with the LDLa, and requires H2 relaxin binding to stabilize an active conformation. Conversely, the GDxxGWxxxF motif in RXFP2 is more closely associated with the LDLa, forming an essential binding interface for H2 relaxin. These differences in the activation mechanism will aid drug development targeting these receptors.

---

The human relaxin family peptide receptors, RXFP1 and RXFP2, are class A G protein-coupled receptors (GPCR) that are activated by the two peptide hormones of the insulin superfamily, H2 relaxin and INSL3 respectively^1^. H2 relaxin is also a high affinity ligand of RXFP2 although the physiological significance of this interaction in humans is unknown. The presence of a Leucine Rich Repeat (LRR) domain within the large N-terminal ectodomain further classifies the receptors as a Leucine-rich repeat containing GPCR (LGR) which includes the glycohormone-binding receptors (GHPRs) and R-spondin receptors (LGR4-6)^2^. An N-terminal low density lipoprotein type A (LDLa) module, connected by a 26- to 32-residue “linker”, further distinguishes RXFP1 and RXFP2 as LGR type C receptors^1, 2^. The LDLa, which requires a structural calcium^3–6^, is indispensable for activation of both RXFP1 and RXFP2^7^, suggesting that the LDLa may be a tethered agonist. Using mutagenesis of the LDLa in whole receptor assays, we showed that residues (Leu7, Lys17) in the LDLa N-terminal region of RXFP1 appeared to be responsible for activity^8, 9^. In contrast, mutation of these residues in the LDLa of RXFP2 had little impact on activity, instead the C-terminal region of the LDLa of RXFP2 may be important for activity^10^. Nevertheless, these activity losses were modest and while the mutations could not pinpoint critical residues/regions that conclusively supported the role of the LDLa as a tethered agonist, they highlighted possible different mechanisms of activation for RXFP1 and RXFP2.

More recently, we showed that the RXFP1 linker comprises a weak-binding site for H2 relaxin that together with the strong-binding site of the LRR domain is essential for ligand-induced activation^11^. Indeed, mutations within the region immediately C-terminal to the LDLa (GDxxGWxxxF, conserved in both RXFP1 and RXFP2) slightly reduced H2 relaxin affinity, but in some cases almost abolished activation^11^. Notably, these critical residues do not appear to directly bind H2 relaxin. Rather, a central transient-helical region of the linker was identified as the ligand-binding site. In NMR experiments, a lack of ^1^H-^1^H NOEs suggested no interaction between linker and the LDLa, so we proposed that H2 relaxin binding to the linker promotes a reorientation of the LDLa to fully engage the activated receptor^11^.

Sequence alignment of RXFP1 and RXFP2 show that the linker of RXFP2 lacks the equivalent H2 relaxin-binding site^12^. Despite this, H2 relaxin binds strongly to RXFP2, through the LRR with only weak contributions to binding from the C-terminal region of the LDLa, including the GDxxGWxxxF motif^12^. Mutations of the conserved residues within the GDxxGWxxxF of RXFP2 induced significant losses of both H2 relaxin binding and subsequent receptor activation^12^. Conversely, INSL3, the cognate ligand for RXFP2, bound normally to the motif-mutant receptor, but receptor activation was completely impaired. Evidently, the linker of these receptors behaves in distinctly different manners depending on both the receptor and ligand involved.

Here we use small angle X-ray scattering (SAXS), NMR spectroscopy, and cell-based receptor assays to show that the LDLa modules and linkers associate differently for the two receptors to form distinct structures. These conformational differences lead to H2 relaxin binding at distinct sites on the LDLa-linker of RXFP1 compared to that of RXFP2. Our results suggest that the conformation of the LDLa-linker and its association with H2 relaxin modulates the structure the GDxxGWxxxF motif, and it is this motif that acts as a tethered agonist, essential for activation of these receptors.

## Results

### The LDLa module is essential for both RXFP1 and RXFP2

At high concentrations H2 relaxin weakly dimerises (K_D_ ∼10 µM) which can be prevented by amidation of the C-termini of H2 relaxin^13^. Therefore, we recombinantly expressed and purified ^15^N-labeled RXFP1 LDLa module and its 32-residue linker (RXFP1_(1-72)_) and performed a 2D ^1^H-^15^N HSQC-monitored titration with amidated H2 relaxin (hereon referred to as relaxin-NH_2_). Compared to previous titrations with H2 relaxin^11^ we observed significantly larger chemical shift changes for similar residues on the linker, Asp51-Thr61 (Fig. 1a, Table 1), with a 3-fold stronger affinity (85 ± 10 μM) than H2 relaxin (200 ± 10 µM)^11^. We also prepared a similar construct of ^15^N-labelled RXFP2 LDLa module and its 26-residue linker (RXFP2_(1-65)_) and titrated it against relaxin-NH_2_. Smaller chemical shift perturbations were observed for residues including Cys26, Asp30 and Glu38, from the C-terminal end of the LDLa, and Asp43-Ile50, from the linker giving an affinity of 205 ± 4 µM (Fig. 1b, Table 1), slightly stronger than previously reported for B5-29 [B-K9R,A-K9/17R] H2 relaxin (330 ± 10 µM)^12^. These findings support the proposal that relaxin has auxiliary binding sites in the LDLa and linker but involves unique binding modes and residues for each receptor.

**Figure 1:**
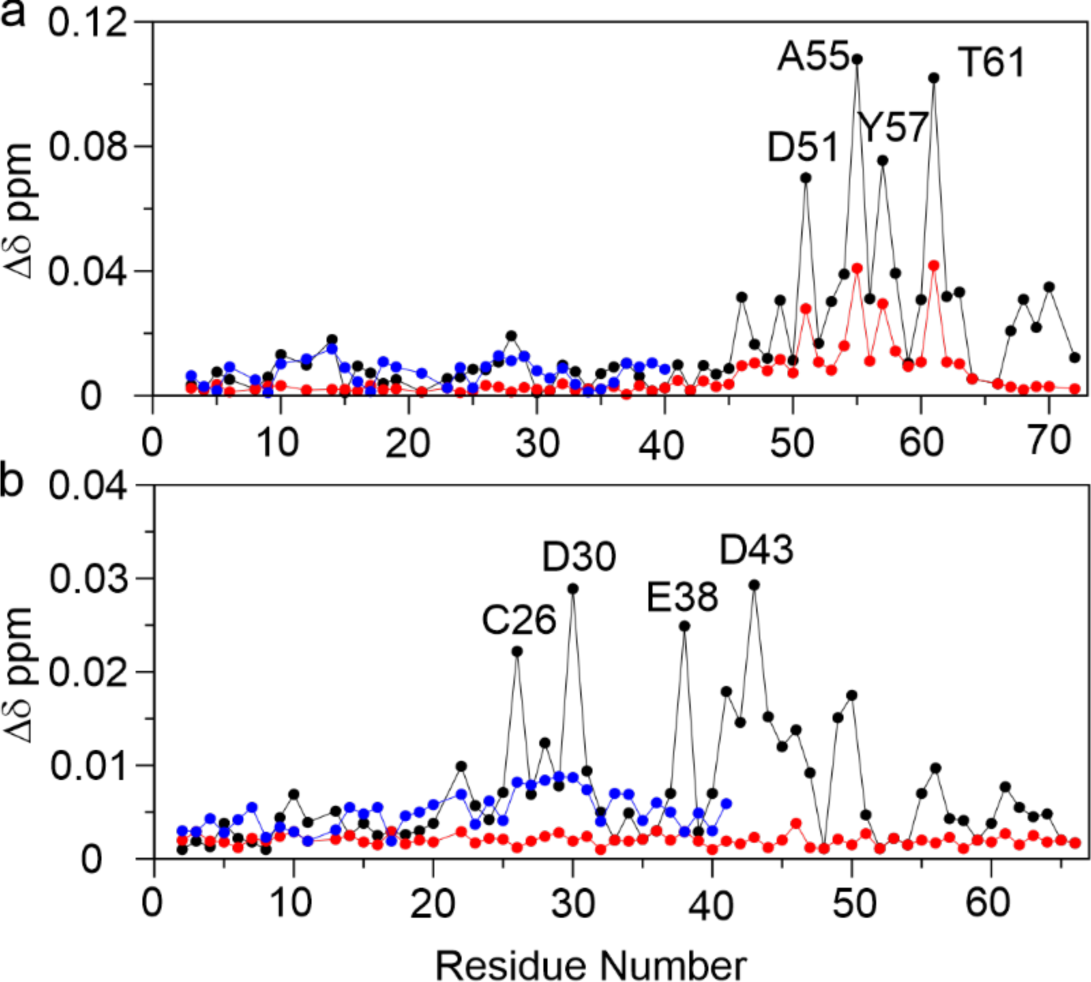
Titrations of RXFP1_(1-72)_ and RXFP2_(1-65)_ with relaxin show different ligand binding sites. Plots of changes in the average ^1^HN and ^15^N chemical shifts differences (Δδ ppm) for titrations with relaxin-NH_2_ of (a) ^15^N-labelled RXFP1_(1–72)_ and (b) ^15^N-labelled RXFP2_(1–65)_ in the presence (black circles) and absence (red circles) of 10 mM CaCl_2_. For comparison in blue circles a similar titration with relaxin-NH_2_ of (a) ^15^N-labelled RXFP1_(1-40)_ and (b) ^15^N-labelled RXFP2_(1-41)_ (with calcium) is shown. Resonances of residues most significantly affected by relaxin-NH_2_ are labelled. Experiments were conducted at pH 6.8 and 25 °C with 25 μM protein and titrated with 20 equivalents of relaxin-NH_2_.

**Table 1:**
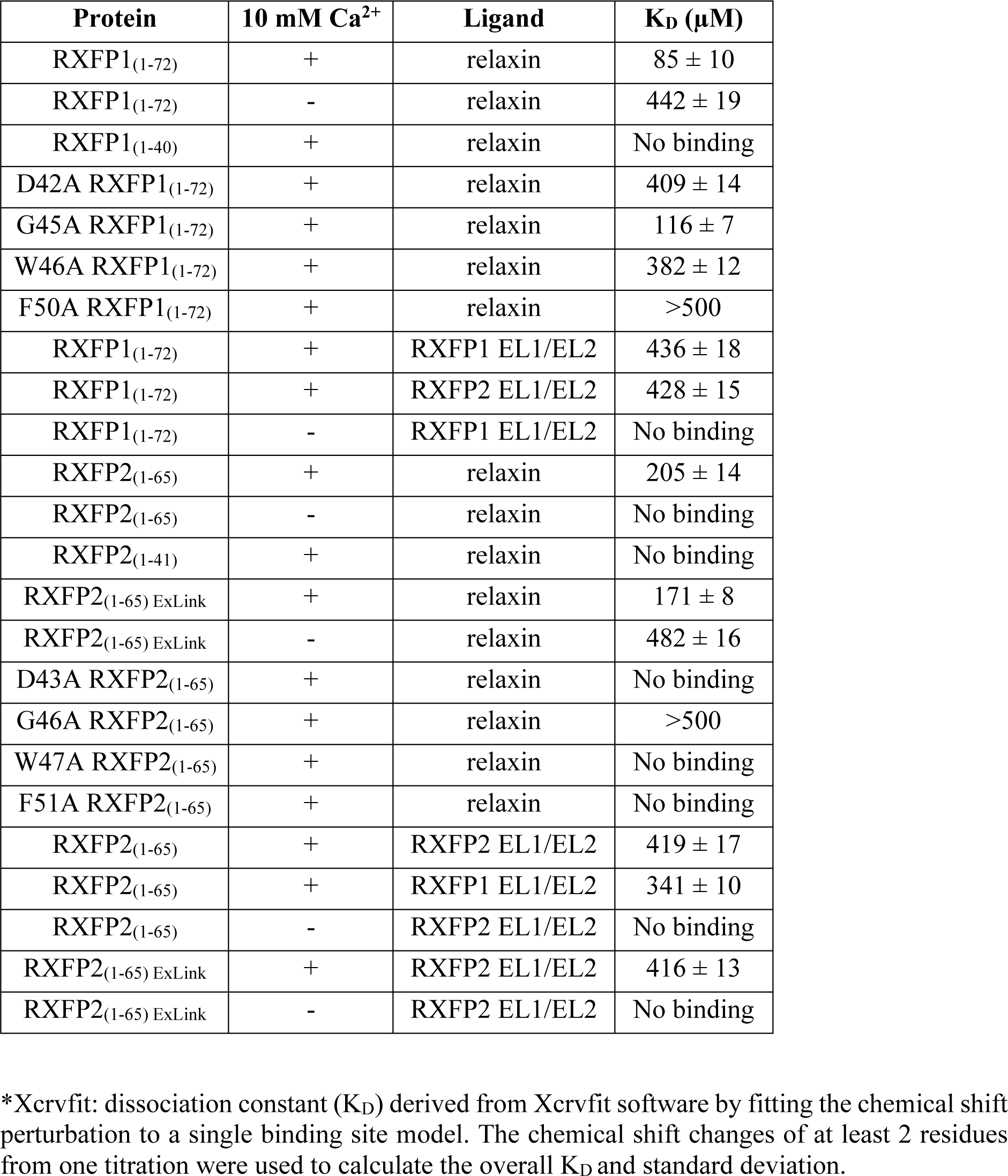
Binding affinities determined by saturation binding (*Xcrvfit*)

| Protein | 10 mM Ca<sup>2+</sup> | Ligand | K<sub>D</sub> (μM) |
| --- | --- | --- | --- |
| RXFP1<sub>(1-72)</sub> | + | relaxin | 85 ± 10 |
| RXFP1<sub>(1-72)</sub> | - | relaxin | 442 ± 19 |
| RXFP1<sub>(1-40)</sub> | + | relaxin | No binding |
| D42A RXFP1<sub>(1-72)</sub> | + | relaxin | 409 ± 14 |
| G45A RXFP1<sub>(1-72)</sub> | + | relaxin | 116 ± 7 |
| W46A RXFP1<sub>(1-72)</sub> | + | relaxin | 382 ± 12 |
| F50A RXFP1<sub>(1-72)</sub> | + | relaxin | >500 |
| RXFP1<sub>(1-72)</sub> | + | RXFP1 EL1/EL2 | 436 ± 18 |
| RXFP1<sub>(1-72)</sub> | + | RXFP2 EL1/EL2 | 428 ± 15 |
| RXFP1<sub>(1-72)</sub> | - | RXFP1 EL1/EL2 | No binding |
| RXFP2<sub>(1-65)</sub> | + | relaxin | 205 ± 14 |
| RXFP2<sub>(1-65)</sub> | - | relaxin | No binding |
| RXFP2<sub>(1-41)</sub> | + | relaxin | No binding |
| RXFP2<sub>(1-65)</sub> ExLink | + | relaxin | 171 ± 8 |
| RXFP2<sub>(1-65)</sub> ExLink | - | relaxin | 482 ± 16 |
| D43A RXFP2<sub>(1-65)</sub> | + | relaxin | No binding |
| G46A RXFP2<sub>(1-65)</sub> | + | relaxin | >500 |
| W47A RXFP2<sub>(1-65)</sub> | + | relaxin | No binding |
| F51A RXFP2<sub>(1-65)</sub> | + | relaxin | No binding |
| RXFP2<sub>(1-65)</sub> | + | RXFP2 EL1/EL2 | 419 ± 17 |
| RXFP2<sub>(1-65)</sub> | + | RXFP1 EL1/EL2 | 341 ± 10 |
| RXFP2<sub>(1-65)</sub> | - | RXFP2 EL1/EL2 | No binding |
| RXFP2<sub>(1-65)</sub> ExLink | + | RXFP2 EL1/EL2 | 416 ± 13 |
| RXFP2<sub>(1-65)</sub> ExLink | - | RXFP2 EL1/EL2 | No binding |

\*Xcrvfit: dissociation constant (K<sub>D</sub>) derived from Xcrvfit software by fitting the chemical shift perturbation to a single binding site model. The chemical shift changes of at least 2 residues from one titration were used to calculate the overall K<sub>D</sub> and standard deviation.

The indispensable role of the LDLa in ligand-mediated activation of both RXFP1 and RXFP2 has been demonstrated^14^. However, the chemical shift changes observed (Fig. 1) on titration with relaxin-NH_2_ suggest that the LDLa in RXFP2 and not RXFP1, has a role in relaxin binding. As the LDLa modules ligate calcium to stabilize their structures^3–6^ we investigated the importance of the folded LDLa in relaxin binding by titrating ^15^N-labelled RXFP1_(1-72)_ and RXFP2_(1-65)_ against relaxin-NH_2_, but in the absence of calcium. For RXFP1_(1-72)_, similar key residues in the linker, Asp51-Thr61, showed chemical shift perturbations, but with a 4-fold decrease in affinity (Fig. 1a, Table 1). For RXFP2_(1-65)_, no significant chemical shift changes in either the LDLa or linker were observed (Fig. 1b, Table 1) suggesting no binding and demonstrating that the fold of the LDLa is essential for binding of relaxin to RXFP2. To further investigate if the folded LDLa alone presents a binding interface for relaxin, we titrated ^15^N-labelled RXFP1_(1-40)_ and RXFP2_(1-41)_ (LDLa module alone without linker) with relaxin-NH_2_ and found that neither showed an interaction (Fig. 1). These data suggest that for RXFP1 the folded LDLa allosterically modulates the affinity of relaxin for the linker; while for RXFP2, the folded LDLa and the linker region Asp43-Ile50 form a structure essential for relaxin binding.

Previously we showed that the LDLa-linker of both RXFP1 and RXFP2 can directly interact with exoloop-1 and -2 (EL1, EL2) of the transmembrane domain (TMD)^11, 12, 15^. These experiments used peptide mimetics of the exoloops, ssRXFP1 (EL1^(475-486)^/EL2-RXFP1) and ssRXFP2 (EL1^(475-486)^/EL2-RXFP2), thus expressing partial exoloop-1 and full-length exoloop-2 with the disulfide that is critical for receptor structure and function^16^. As these experiments were conducted in the presence of calcium, hence a structured LDLa, we tested the importance of a folded LDLa module, by titrating ^15^N-labelled RXFP1_(1-72)_ and RXFP2_(1-65)_, in the absence of calcium, with ssRXFP1 and ssRXFP2 and observed a complete lack of interaction (Supplementary Fig. 1). These data suggest that while the linker region contains all the key residues, it requires structured LDLa modules to effectively interact with H2 relaxin and the TMD. Hence for both receptors, RXFP1 and RXFP2, it appears that the LDLa serves as an anchor that structures the linker region and this interaction is critical for receptor activation induced by the binding to H2 relaxin^11, 17^.

### The LDLa-LRR linkers of RXFP1 and RXFP2 adopt distinct conformational ensembles

To understand the structural relationship between the LDLa and the linker, we performed SAXS on RXFP1_(1-72)_ and RXFP2_(1-65)_ (Fig. 2a). For RXFP1_(1-72),_ the radius of gyration (*R*_g_) was 21.06 ± 0.19 Å and the molecular weight as 7.68 kDa (expected M.W. ∼8.29 kDa) (Fig. 2b,c and Supplementary Table 2). For RXFP2_(1-65)_, *R*_g_ and molecular weight were measured to be 15.98 ± 0.14 Å and 6.35 kDa (expected M.W. ∼7.09 kDa) respectively. The shapes of RXFP1_(1-72)_ and RXFP2_(1-65)_ were determined *ab initio* using DAMMIN (Supplementary Fig. 2). To further model different possible conformations of RXFP1_(1-72)_ and RXFP2_(1-65)_ in solution, we used the Ensemble Optimization Method (EOM)^18, 19^ to generate a pool of different conformations (Fig. 2d). The range of conformations for RXFP1_(1-72)_ is smaller compared to RXFP2_(1-65)_, which shows two distinct pools. Collectively, the SAXS models of RXFP1_(1-72)_ suggest that the linker folds back onto the LDLa, providing a more globular shape, whereas for RXFP2_(1-65)_, the models suggest an elongated linker extending away from the LDLa.

**Figure 2:**
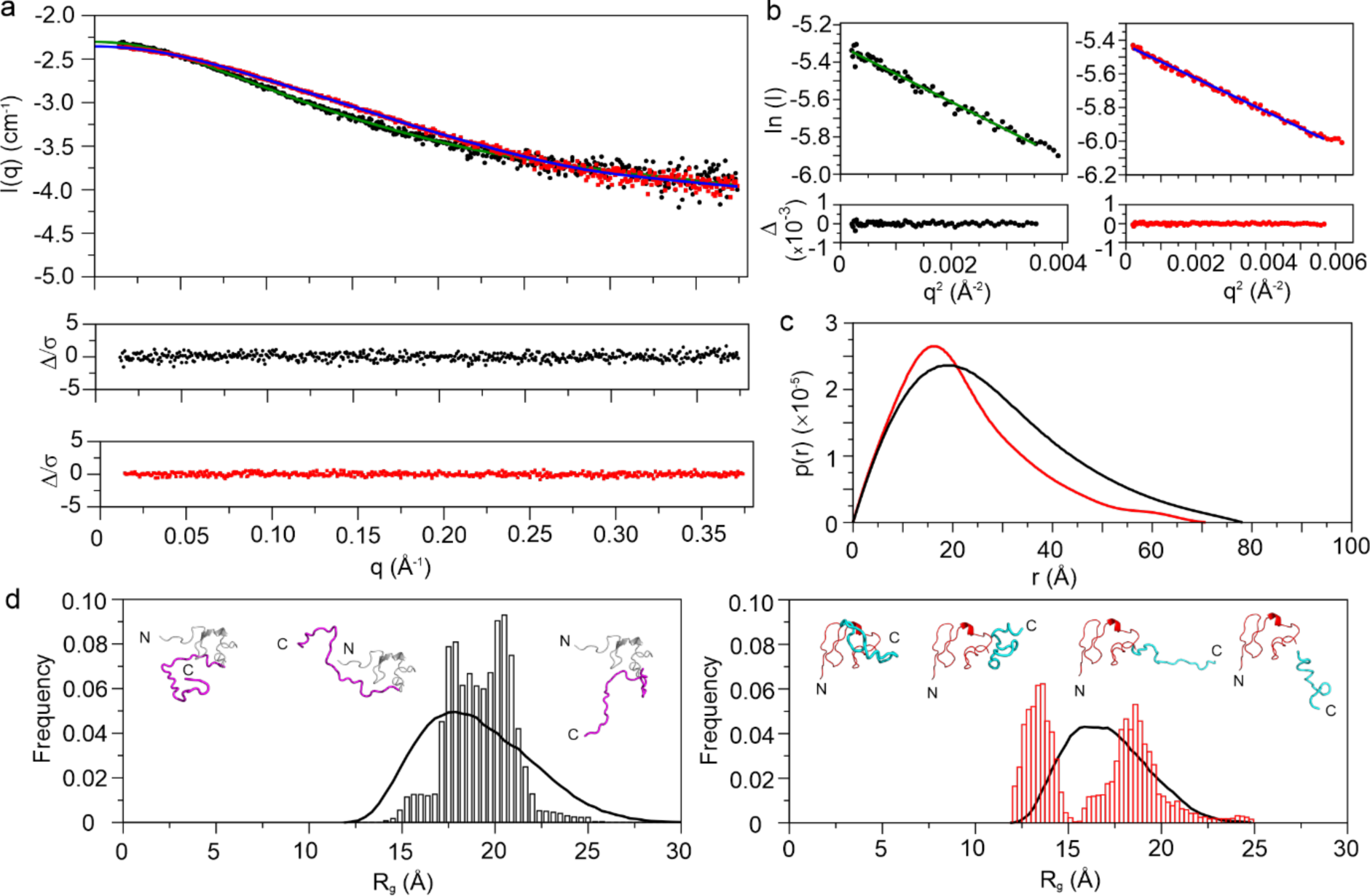
Small Angle X-Ray Scattering analysis shows the linker and LDLa modules interact differently for RXFP1_(1-72)_ and RXFP2_(1-65)_. (a) Small Angle X-Ray Scattering data for RXFP1_(1-72)_ (black) and RXFP2_(1-65)_ (red). (b) Guinier Approximation curve for RXFP1_(1-72)_ (black) and RXFP2_(1-65)_ (red). (c) p(r) curve for RXFP1_(1-72)_ (black) and RXFP2_(1-65)_ (red) highlighting the difference in their radius of gyration. (d) EOM modeling distribution along with the models proposed for RXFP1_(1-72)_ (left, LDLa module in grey, linker in magenta) and RXFP2_(1-65)_ (right, LDLa module in red, linker in cyan) showing a bimodal distribution for RXFP2_(1-65)_.

Due to lack of ^1^H-^1^H NOEs between the linker and LDLa, we could not solve the structure of the LDLa-linker by NMR. We collected ^13^Cαβ chemical shift and ^15^N{^1^H}-NOE data for RXFP1_(1-72)_ in the presence of relaxin-NH_2_ and compared to data in the absence of relaxin-NH_2_ (^11^ and Fig. 3a). As expected, the ΔCα-ΔCβ and ^15^N{^1^H}-NOE data suggest a slight increase in the extent and stability of helix for the region Leu48-Tyr58 than previously observed for H2 relaxin^11^ due to stronger affinity and the monomeric nature of relaxin-NH_2_. For RXFP2_(1-65)_ (Fig. 3b), we observed ΔCα-ΔCβ values consistent with the secondary structure of the RXFP2 LDLa^10^. In the absence of relaxin-NH_2_, the ΔCα-ΔCβ values suggest that the RXFP2 linker region is largely unstructured, except for the Gly42-His55 that showed negative ΔCα-ΔCβ values indicating the presence of transient β-structure. To investigate this structure, we also recorded ^15^N{^1^H}-NOEs on ^15^N-labelled RXFP2_(1-65)_. The average ^15^N{^1^H}-NOE values (0.76 ± 0.10) (Supplementary Table 1) for the LDLa (Ser5-Cys41) agree with a folded structure (PDB 2m96). Notably, residues in the RXFP2-linker from Gly42-Phe51 showed ^15^N{^1^H}-NOE values ranging from 0.49 to 0.84 (average 0.62 ± 0.10), consistent with transient structure whereas the corresponding region in RXFP1_(1-72)_ (Gly41-Phe50) is more flexible showing a range of ^15^N{^1^H}-NOE values of 0.35 to 0.70 (average 0.53 ± 0.09) (Fig. 3a; Supplementary Table 1). For RXFP2_(1-65)_ following Phe51, the ^15^N{^1^H}-NOE value progressively decreases, implying a flexible C-terminal region extending away from the LDLa. These ^15^N{^1^H}-NOE data complement the SAXS EOM analysis for both RXFP1_(1-72)_ and RXFP2_(1-65)_, and further suggest that the linker region immediately C-terminal to the LDLa in RXFP2_(1-65)_ is more ordered than in RXFP1_(1-72)_. In contrast to RXFP1_(1-72)_, on addition of relaxin-NH_2_ to ^15^N-labelled RXFP2_(1-65)_, little change is observed in the ΔCα-ΔCβ and ^15^N{^1^H}-NOE profiles, indicating no significant structural change (Fig. 3).

**Figure 3:**
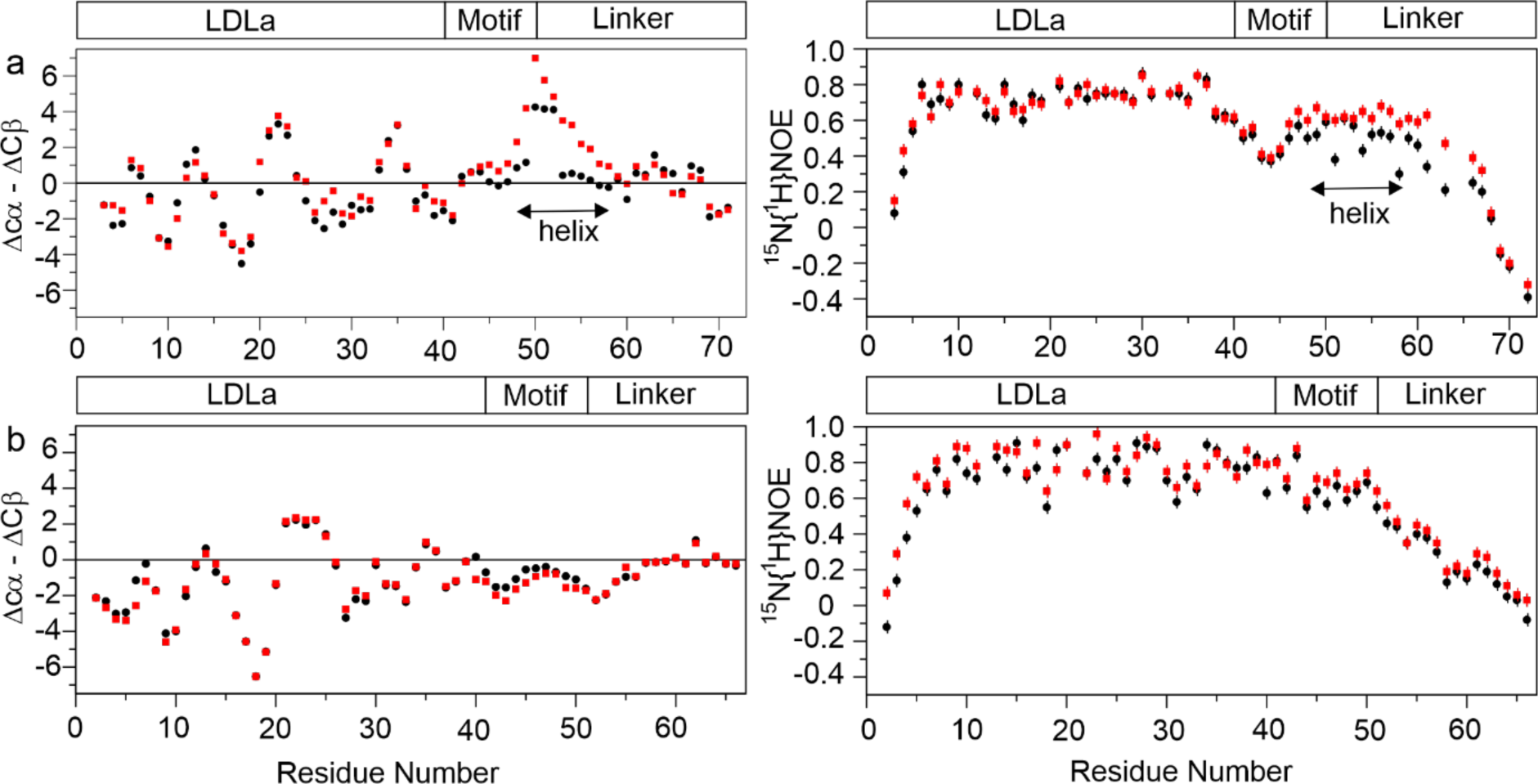
Interaction with relaxin shows conformational changes for RXFP1_(1-72)_ but not RXFP2_(1-65)_. Plots of ^13^Cαβ secondary chemical shifts (left panels) and ^15^N^29^-NOEs (right panels) for apo- (black circle) and in the presence of 20 equivalents of amidated relaxin (red square) for (a) RXFP1_(1-72)_ and (b) RXFP2_(1-65)_. The regions encompassing the LDLa module the GDxxGWxxxF motif and the remainder of the linker for both receptors are indicated as a block diagram. A distinct increase in stabilization of helical structure is observed for RXFP1_(1-72)_ for residues within the linker (Leu48-Tyr58) whereas addition of relaxin to RXFP2_(1-65)_ shows little change to structure despite clear binding (Figure 1).

### Structural effect of mutations in the GDxxGWxxxF motif of RXFP1 and RXFP2

The conserved residues within the GDxxGWxxxF motif are critical for H2 relaxin binding and activation of RXFP1 and RXFP2^11, 12^. Mutation of Asp42, Gly45, Trp46 and Phe50 from this motif to Ala in RXFP1 reduced H2 relaxin-induced cAMP production and weakened the affinity of H2 relaxin to the mutant RXFP1^11^. Equivalent mutations in RXFP2 (Asp43, Gly46, Trp47 and Phe51) caused an even more profound result, completely preventing binding of H2 relaxin and abolishing cAMP production^12^. These mutations did not affect the binding of the cognate-ligand INSL3 to RXFP2, but reduced or abolished INSL3 activation of the receptor^12^. Here we translated these single residue mutations of the motif into our recombinant constructs RXFP1_(1-72)_ and RXFP2_(1-65)_ and characterized the impact of these mutations on structure by monitoring ^1^H,^15^N chemical shifts, ^15^N{^1^H}-NOE and relaxin-NH_2_ binding (Fig. 4, Table 1).

**Figure 4:**
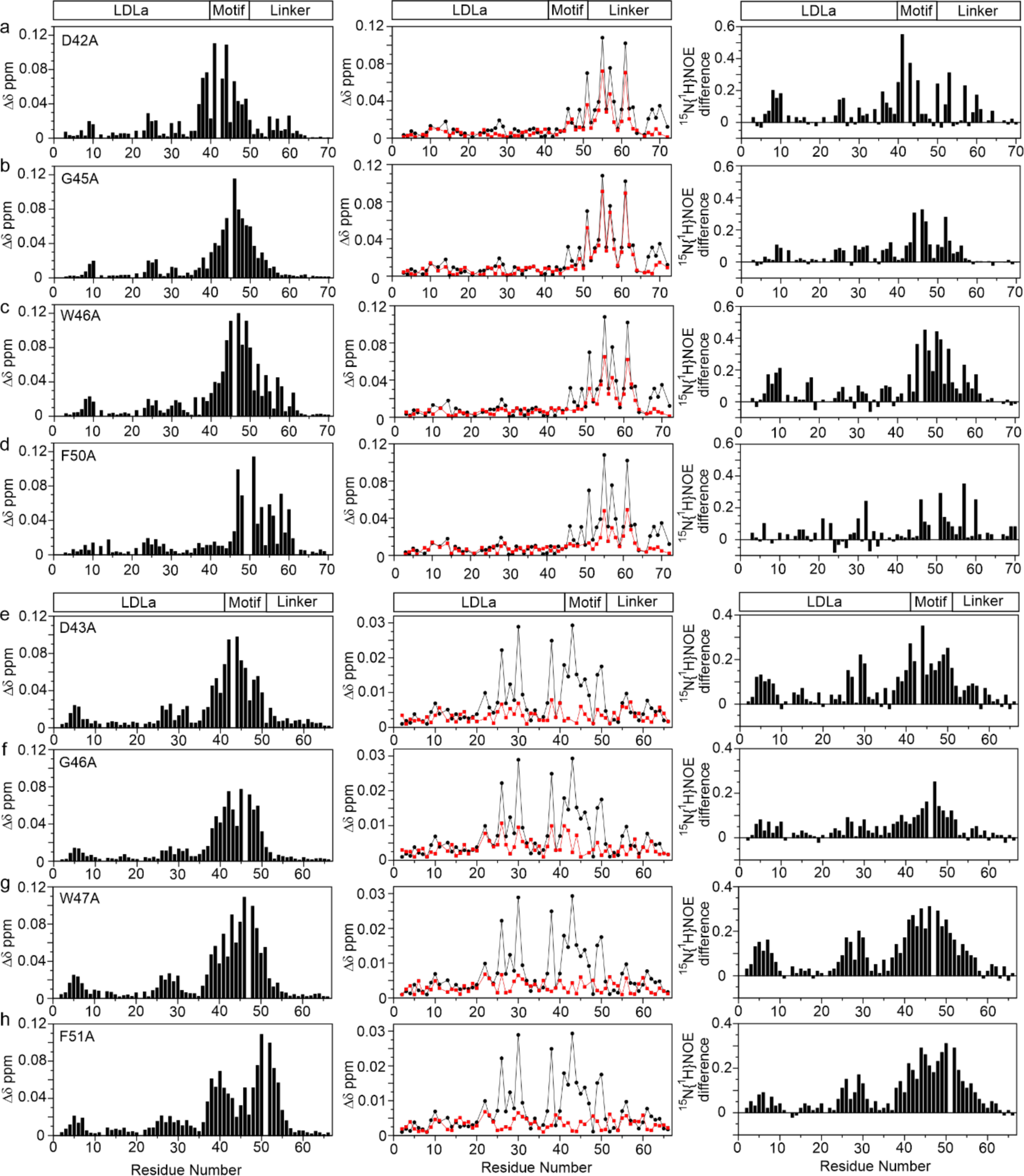
Mutations within the GDxxGWxxxF motif have different effects on structure and relaxin binding of RXFP1_(1-72)_ and RXFP2_(1-65)_. For (a-h) the first column is the average chemical shift changes of the ^1^HN and ^15^N (Δδ ppm) for mutant compared to wild-type; the second column shows the average chemical shift differences the ^1^HN and ^15^N (Δδ ppm) after titration with 20 equivalents of relaxin for mutant (red squares) and wild-type (black circles); the third column is the difference (wild-type less mutant) of the ^15^N{^1^H}-NOE where a positive difference indicates a lower NOE in the mutant. The following mutations were made for RXFP1_(1-72)_ (a) (D42A) (b) G45A (c) W46A (d) F50A and for RXFP2_(1-65)_ (e) D43A (f) G46A (g) W47A (h) F51A. A block diagram above (a) and (e) show the sequence location of the LDLa module, the GDxxGWxxxF motif and the remainder of the linker in RXFP1 and RXFP2.

Upon titration of D42A, G45A and W46A-RXFP1_(1-72)_ with relaxin-NH_2_, both D42A and W46A showed a ∼4-fold decrease in affinity of relaxin-NH_2_ to the linker (Table 1). In comparison to wild-type RXFP1_(1-72)_, D42A showed extensive amide chemical shift perturbation and a decrease in the ^15^N{^1^H}-NOE values for the linker region, Gly41-Phe50 and to a lesser degree for amide resonances up to Met60 (Fig. 4a). W46A had a significant impact on the amide chemical shifts and ^15^N{^1^H}-NOE values for the residues that form the helical structure (Leu48-Thr61) and hence affected relaxin-NH_2_ binding (Fig. 4c). Titration of F50A with relaxin-NH_2_ showed >6-fold loss of ligand binding (Table 1). F50A showed large chemical shift perturbations and a decrease in the ^15^N{^1^H}-NOE values for Asp51-Thr61, a region which is essential for ligand binding (^11^ and Fig. 4d). In contrast, G45A had minimal effect on relaxin-NH_2_ binding (Table 1) but showed similar chemical shift and ^15^N{^1^H}-NOE perturbations to W46A (Fig. 4b), although not as profound. Importantly, for all mutants we observed subtle changes in the chemical shifts and the ^15^N{^1^H}-NOE values of resonances of the LDLa. For D42A and W46A consistent changes were seen in the N-terminal region of the LDLa, Leu7-Gly10 and those immediately after the 3_10_-helix, His24-Asn26, which supports the linker folding around the LDLa, as suggested by the SAXS EOM analysis (Fig. 2d).

For RXFP2_(1-65)_, the equivalent mutations, D43A, W47A and F51A, showed near complete loss of binding to relaxin-NH_2_ and for G46A, significantly reduced binding (Fig. 4e-h, Table 1). Importantly, the mutations had widespread effect both on the amide chemical shifts and ^15^N{^1^H}-NOE values. Similar to RXFP1, we observed chemical shift perturbations and a decrease in the ^15^N{^1^H}-NOE values for the residues from the N-terminus of the LDLa, Ile2-Tyr10, but also to residues encompassing Cys26-Asp30, but not the 3_10_-helix prior to Cys26. Notably, Cys26-Asp30 is the same region shown to participate in binding to relaxin-NH_2_ (Fig. 1b). Together, the data suggest that mutations in the motif GDxxGWxxxF structurally impact specific regions of the LDLa modules supporting interactions between the modules and linkers of the two receptors.

### Functional effect of swapping the GDxxGWxxxF motif between receptors

The LDLa modules of RXFP1 and RXFP2 can be swapped with limited effect on H2 relaxin binding and ligand-induced activation of these chimeras^20^. For example, the chimera RXFP211, comprising the LDLa (Met1-Gly42) of RXFP2 with the linker, LRR domain and TMD of RXFP1, shows compared to RXFP1, similar affinity to H2 relaxin but with slightly lower efficacy in ligand-stimulated cAMP activity compared to RXFP1 (^20^ and Fig. 5a). To compare the role of the conserved GDxxGWxxxF motif in both receptors, we designed a chimera (RXFP2F11) made up of the LDLa and GDxxGWxxxF motif (Met1-Phe51) of RXFP2 and the remainder of RXFP1. This chimera expressed at the cell surface at a slightly higher level than RXFP1 and similar to the level of RXFP211 (Fig. 5c). However, compared to these receptors, RXFP2F11 had a greatly reduced signalling capacity in response to H2 relaxin (Fig. 5a,d), and showed minimal binding (K_D_ >20 nM) of Europium-labelled H2 relaxin at concentrations up to 10 nM (Fig. 5b). These data suggest that swapping the RXFP2 LDLa including the GDxxGWxxxF motif markedly weakens ligand affinity pointing to critical structural differences in how this motif relates to the receptor to modulate H2 relaxin binding and activation.

**Figure 5:**
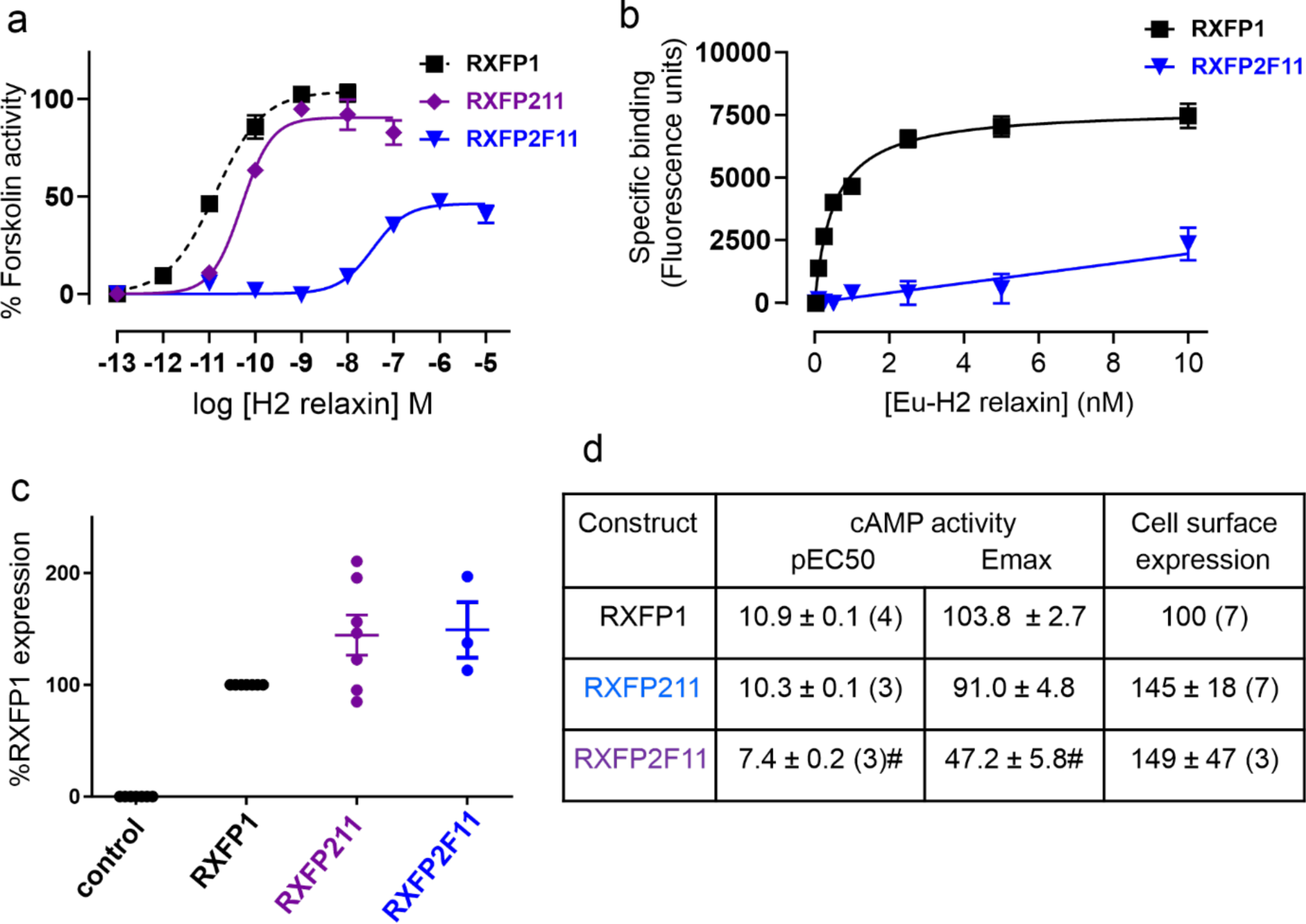
Activity of RXFP1 LDLa-linker chimeric receptors compared to RXFP1. (a) H2 relaxin-induced cAMP responses expressed as % 5 µM Forskolin response (b) Eu-H2 relaxin saturation binding where RXFP1 shows an affinity (K_D_) of 0.6 nM and RXFP2F11 an affinity of > 20 nM (the binding affinities reflect the sensitivity of the assay, maximum H2 concentration 15 nM). (c) Cell surface expression of chimeric receptors compared to RXFP1. Data are expressed as mean ± S.E.M of triplicate determinations from at least 3-7 independent experiments. (d) Pooled cAMP activity (pEC50 and Emax) and cell surface expression (%RXFP1) data for RXFP211 and RXFP2F11 in comparison to RXFP1. #p<0.01 compared to RXFP1.

We have previously shown H2 relaxin binding to RXFP1 is biphasic and to RXFP2 is monophasic, highlighting mechanistic differences^17^. Sequence alignments show there is a 7-residue insertion in the linker of RXFP1 that defines a part of its relaxin-binding site. By inserting these residues (Lys52-Tyr58) into the linker of RXFP2 we recapitulated the biphasic binding and dissociation of H2 relaxin^17^. This chimera, in response to H2 relaxin activation, showed greater potency and efficacy than RXFP2 and less potency, but similar efficacy to RXFP1. We inserted these seven residues into RXFP2_(1-65)_ (RXFP2_(1-65)_-ExLink2), ^15^N-labelled it and showed in the presence of calcium the affinity to relaxin-NH_2_ was 2-fold weaker than RXFP1_(1-72)_, but 3.5-fold stronger than RXFP2_(1-65)_ (171 µM, Table 1). In the absence of calcium, the affinity weakened (482 µM) to be similar to RXFP1_(1-72)_ (442 µM). As expected, these titrations showed a change in the relaxin-NH_2_ binding site to the extended linker region, similar to that of RXFP1_(1-72)_ (Supplementary Fig. 2a, Fig. 1). Further, titration of ^15^N-labelled RXFP2_(1-65)_-ExLink2 with the TMD exoloop mimetic ssRXFP2, showed this mimetic also interacted within the extended linker, similar to RXFP1_(1-72)_ and with comparable affinity (Table 1, Supplementary Fig. 1c).

To test whether the 7-residue insertion into RXFP2_(1-65)_ influence or is influenced by the GDxxGWxxxF motif, we investigated the structural properties of the RXFP2_(1-65)_-ExLink2. ^13^Cαβ secondary shifts and ^15^N{^1^H}-NOE data showed that extending the linker region surprisingly did not induce helical propensity in either apo- or relaxin-bound states (Supplementary Fig. 2b,c). The average ^15^N{^1^H}-NOE value for the residues Lys53-Glu73 that are C-terminal to the GDxxGWxxxF motif was 0.17 ± 0.08 which supports a lack of structural ordering. The average ^15^N{^1^H}-NOE value for the GDxxGWxxxF motif in RXFP2_(1-65)_-ExLink2 (0.63 ± 0.15) was comparable to that recorded for RXFP2_(1-65)_ (0.64 ± 0.10) suggesting no change in the dynamic properties of the motif. These data suggest that transferring the RXFP1 linker-relaxin binding site into the RXFP2 linker changes the ligand binding interface and affinity but does not allow the formation of helical structure and/or stabilization post-ligand binding, as observed for RXFP1. These data collectively demonstrate that the fold of the motif GDxxGWxxxF in RXFP1 and RXFP2 modulates the structuring of the linker and its association with the LDLa, explaining why swapping the motif between receptors (such as RXFP2F11) led to significant loss of H2 relaxin binding and potency.

### Fast timescale dynamics of apo- and relaxin-bound RXFP1_(1-72)_ and RXFP2_(1-65)_

The differences in conformation and relaxin-binding properties of the LDLa and linker of RXFP1_(1-72)_ and RXFP2_(1-65)_ suggested differences in dynamics. To investigate this possibility, we acquired ^15^N spin relaxation experiments, ^15^N-R_1_, ^15^N-R_2_ and ^15^N{^1^H}-NOE at 81.1 MHz for RXFP1_(1-72)_ and RXFP2_(1-65)_ in the presence or absence of relaxin-NH_2_ and at 25 or 15 °C (Fig. 6). In addition, the data were subjected to a reduced spectral density analysis (supplementary Fig. 3). For RXFP1_(1-72)_ the spin-relaxation values (Fig. 6) show that the first six and last ten residues are disordered and insensitive to relaxin-NH_2_, so we focused our analysis on Ser6-Cys40 of the LDLa, and Gly41-Met60 of the linker. Similarly, the first four residues of the LDLa of RXFP2_(1-65)_ and residues after Gly52 are essentially disordered. Therefore, for RXFP2_(1-65)_ we focused on Ser5-Cys41 of the LDLa and Gly42-Gly52 of the linker. The average values for ^15^N-R_1_, ^15^N-R_2_ and ^15^N{^1^H}-NOE, and the reduced spectral parameters J(0), J(0.87ω_H_) and J(ω_N_) for these regions are provided in Supplementary Table 1.

**Figure 6.**
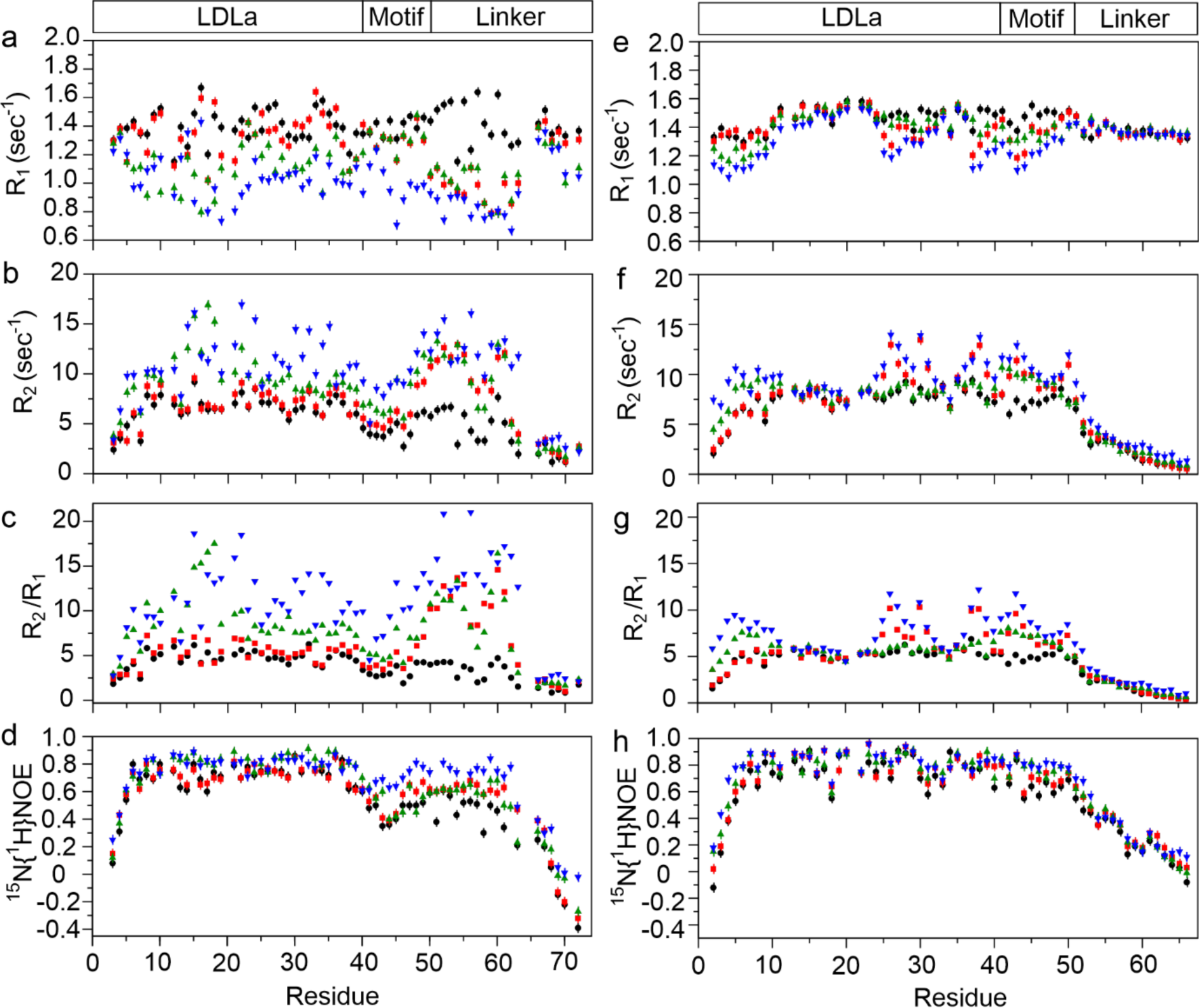
^15^N-spin relaxation dynamics of RXFP_(1-72)_ and RXFP2_(1-65)_. Backbone amide ^15^N relaxation parameters of 100 µM RXFP1_(1-72)_ (a-d) and RXFP2 _(1-65)_ (e-h) in apo-state at 25 °C (black circles), and at 15 °C (green triangle) and bound with amidated relaxin at 25 °C (red circle) and at 15 °C (blue triangle) measured at ^15^N frequency of 81.1 MHz and pH 6.8. Sample concentrations represent 83% saturation of RXFP1_(1-72)_ and RXFP2_(1-65)_ with ligand. Error bars are calculated using Monte Carlo Simulations for ^15^N-R_1_, ^15^N-R_2_ measurements and based on average estimated noise level for ^15^N{^1^H}-NOE. The sequence locations of the LDLa module, the GDxxGWxxxF motif and the remainder of the linker in the receptors are shown above (a) and (e).

For RXFP1_(1-72)_ the ^15^N{^1^H}-NOE values change little for the LDLa under all conditions, consistent with a well-ordered structure. In contrast, ^15^N{^1^H}-NOE values for the linker of RXFP1_(1-72)_ increase at 15 °C or on addition of relaxin-NH_2_ suggesting a gain in order. As expected, at 25 °C and in the presence of relaxin-NH_2_, the ^15^N-R_2_ for residues of the linker increase, especially Gln49-Met60, which is clearly observed in the R_2_/R_1_ ratios (Fig. 6), reflecting the fast-timescale interaction with relaxin-NH_2_. However, at 15 °C and in the *absence* of relaxin-NH_2_, there are marked increases in R_2_/R_1_ (>10) for residues in the relaxin-binding site, Phe50-Ala55, Lys59-Thr61, supporting the presence of transient structure. Notably, there are similar increases in R_2_/R_1_ (>10) for residues within the LDLa, Gly8, Phe10, Cys12, and Asn14-Cys18 (Fig. 6b). These pronounced changes pose the question: are the dynamics for these residues of the LDLa connected to those of the linker? To partly address this question, we collected ^15^N spin relaxation data for the LDLa alone at 15 and 25 °C (Supplementary Fig. 4). The only residue to show a significant R_2_/R_1_ (>10) was Lys17 at 15 °C. We investigated the effect of decrease in temperature (15 °C) on the dynamics of RXFP1_(1-72)_ bound to relaxin-NH_2_. With the exception of the GDxxGWxxxF motif, the LDLa and linker show similar average ^15^N{^1^H}-NOE and ^15^N-R_2_ values (Fig. 6d, Supplementary Table 1). However, significant R_2_/R_1_ (>10) are observed for the entire linker from Gly45 to Gln63 and for extensive regions of the LDLa, Ser6, Cys12, Asn14, Ile15, Lys17-His24, Gly27, Val28, Asp30, Gly32, Gln34, Ala35 and Asp38. Importantly, the reduced spectral density parameters J(0) and J(0.87ω_H_) for the LDLa and linker were now similar, reflecting similar ordering despite the distinct chemical exchange. Furthermore, on addition of relaxin-NH_2_, J(0) increased and J(0.87ω_H_) decreased for the linker (Supplementary Table 1), consistent with an increase in ordering within the linker.

For RXFP2_(1-65)_ under all conditions, the ^15^N{^1^H}-NOE over Gly5-Gly52, covering the LDLa and a substantial part of the linker, are similar, reflecting an overall similar tumbling and ordering of these residues with little conformational change, consistent with the SAXS analysis (Fig. 2), and distinctly different to RXFP1_(1-72)_. Addition of relaxin-NH_2_ at 15 °C shows increased ^15^N-R_2_ and decreased ^15^N-R_1_, clearly observed in the increased R_2_/R_1_, although notably these differences are substantially less than those of RXFP1_(1-72)_. Three regions show distinct changes in R_2_/R_1_ (>8): the N-terminal of the LDLa, Phe4-Lys8; a region within the LDLa, His25-Asp31; and the C-terminal of the LDLa that includes Cys41-Gly46, and Ile50 of the linker. At 15 °C in the absence of relaxin-NH_2_ (Fig. 6c), we observed small but distinct increases in R_2_/R_1_ for residues Ile2-Tyr10 and Glu38-Ile50. These changes are consistent with the reported spatial proximity of N- and C-terminal regions^12^ and so the change in R_2_/R_1_ in the presence of relaxin-NH_2_ for Ile2-Tyr10 may not reflect a direct interaction with relaxin-NH_2_. The remaining two regions (His25-Asp31 and Cys41-Gly46), however, correlate well with the chemical shift differences observed upon relaxin-NH_2_ binding (Fig. 1), and therefore we propose that these increases in R_2_/R_1_ reflect chemical exchange from an interaction with relaxin-NH_2_ and in contrast to RXFP1_(1-72)_, not a conformational change. The reduced spectral density parameters J(0) and J(0.87ω_H_) for the LDLa (Ser5-Cys41) and linker (Gly42-Gly52) of RXFP2_(1-65)_ reflect similar ordering and structure (Supplementary Table 1). Furthermore, on addition of relaxin-NH_2_ at 15 °C, the more modest increases in J(0) and decreases in J(0.87ω_H_) for the LDLa and linker are consistent with the notion of little conformational change in RXFP_(1-65)_.

### 15N-relaxation dispersion on RXFP1_(1-72)_ and RXFP2_(1-65)_

To further understand the influence of relaxin binding on the dynamics of RXFP1_(1-72)_ we acquired ^15^N-CPMG relaxation dispersion experiments in the absence and presence of relaxin-NH_2_ at 15 °C at 70.9 and 81.1 MHz (Table 2; Supplementary Fig. 5a,b). In the absence of relaxin-NH_2_ four residues, Cys12 and Thr16 from the LDLa and Tyr53 and Ser56 from the linker, showed distinct dispersion fitting to a slow-limit exchange, two-state model. As Thr53 and Ser56 are within the transient helix they likely reflect its fluctuations; whereas Cys12 and Thr16 are located in the loop between the β-strands of the LDLa. Both pairs of residues show a population of 2-3% for the minor state. The kinetics vary for the two sites where the linker residues show a k_ex_ of 800 s^-1^ whereas the LDLa residues, a k_ex_ of 250 s^-1^. The kinetics of the linker residues may reflect both the formation of transient structure and interaction with the LDLa, whereas the kinetics of the LDLa residues may only reflect an interaction with a structured linker.

**Table 2:** ^15^N CPMG relaxation dispersion kinetics parameters for ^15^N RXFP1_(1-72)_ and ^15^N RXFP2_(1-65)_

| $^{15}\text{N}$ RXFP1<sub>(1-72)</sub> | | | | | | | | | |
| --- | --- | --- | --- | --- | --- | --- | --- | --- | --- |
| 15 °C relaxin bound |  |  |  |  | 15 °C Apo |  |  |  |  |
| Residue | $k_{\text{ex}}$ ($\text{s}^{-1}$) | pB (%) | $k_{\text{on}}$ ($\times 10^6$) $\text{M}^{-1} \text{s}^{-1}$ | $k_{\text{off}}$ ($\text{s}^{-1}$) | Residue | $k_{\text{ex}}$ ($\text{s}^{-1}$) | pB (%) | $k_{\text{on}}$ ($\times 10^6$) $\text{M}^{-1} \text{s}^{-1}$ | $k_{\text{off}}$ ($\text{s}^{-1}$) |
| **Linker** |  |  |  |  | **Linker** |  |  |  |  |
| Trp46 | 775±29 | 2.9±0.2 | 23±2 | 752±30 | | | | | |
| Tyr53 | 765±53 | 4.3±0.7 | 33±2 | 731±39 | Tyr53 | 845±71 | 1.9±0.7 | 16±2 | 829±78 |
| Ser56 | 551±38 | 5±0.4 | 28±9 | 523±61 | Ser56 | 743±60 | 2.2±0.6 | 17±4 | 727±68 |
| Met60 | 673±55 | 2.7±0.8 | 18±5 | 655±52 | | | | | |
| **LDLa module** |  |  |  |  | **LDLa module** |  |  |  |  |
| Tyr9 | 77±11 | 6.1±0.2 | 4.6±1.1 | 72±8 | | | | | |
| Phe10 | 81±10 | 9.2±0.6 | 7.5±0.9 | 74±9 | | | | | |
| Cys12 | 71±11 | 7.7±0.1 | 5.5±0.9 | 66±5 | Cys12 | 298±41 | 2.4±0.3 | 7.2±0.7 | 291±24 |
| Thr16 | 65±6 | 9.0±0.5 | 5.9±0.8 | 60±6 | Thr16 | 212±33 | 2.5±0.6 | 5.3±1.1 | 207±31 |
| Leu22 | 51±7 | 6.9±0.8 | 3.6±0.8 | 47±8 | | | | | |
| Leu23 | 71±5 | 6.4±0.3 | 4.5±1.1 | 66±4 | | | | | |
| **L7K17 mutant** |  |  |  |  | **L7K17 mutant** |  |  |  |  |
| Tyr53 | 519±20 | 3.4±0.7 | 17.7±1.2 | 501±28 | Tyr53 | 941± 49 | 2.3±0.2 | 22±2 | 919±49 |
| Ser56 | 797±29 | 3.7±0.4 | 29.1±1.9 | 768±29 | | | | | |
| Met60 | 725±41 | 2.4±0.2 | 17.2±1.2 | 707±41 | | | | | |
| $^{15}\text{N}$ RXFP2<sub>(1-65)</sub> | | | | | | | | | |
| 15 °C relaxin bound |  |  |  |  | 15 °C Apo |  |  |  |  |
| Residue | $k_{\text{ex}}$ ($\text{s}^{-1}$) | pB (%) | $k_{\text{on}}$ ($\times 10^6$) $\text{M}^{-1} \text{s}^{-1}$ | $k_{\text{off}}$ ($\text{s}^{-1}$) | Residue | $k_{\text{ex}}$ ($\text{s}^{-1}$) | pB (%) | $k_{\text{on}}$ ($\times 10^6$) $\text{M}^{-1} \text{s}^{-1}$ | $k_{\text{off}}$ ($\text{s}^{-1}$) |
| **Linker** |  |  |  |  | **Linker** |  |  |  |  |
| Gly42 | 30±10 | 37±2 | 11±3 | 19±4 | Gly-42 | 71±20 | 11±2 | 11±2 | 60±8 |
| Trp47 | 52±7 | 30±1 | 16±1 | 37±7 | | | | | |
| **LDLa module** |  |  |  |  | **LDLa module** |  |  |  |  |
| Ser5 | 52±7 | 32±1 | 17±1 | 35±9 | Ser-5 | 112±21 | 8±1 | 9±1 | 103±17 |
| His25 | 71±10 | 26±3 | 19±2 | 52±10 | | | | | |

Upon addition of relaxin-NH_2_, Tyr53 and Ser56 from the linker continue to show chemical exchange, but also Trp46 and Met60. These residues show similar kinetics as viewed in the apo-state with a minor-state population of 2-5% and a k_ex_ of 700 s^-1^. The similar kinetics is surprising, but exchange rates now will reflect a combination of stabilizing transient structure, interactions with both the LDLa and non-saturating relaxin-NH_2_. Within the LDLa, Cys12 and Thr16, but also Tyr9, Phe10, Leu22 and Leu23 show dispersion. These residues all show a significantly larger minor-state population of 6-9% and a 4-fold lower k_ex_ of 65 s^-1^. As previous experiments^11^ and the data presented here show that the LDLa does not interact with relaxin, we interpret these kinetics for the LDLa residues as reflecting only an interaction with the linker, presumably when structured, which is more populated in the presence of relaxin-NH_2_ compared to the apo-state.

Previously we showed that mutation of several residues within the LDLa, mainly L7K and K17A, decreased ligand-binding induced cAMP production which was interpreted to be important for interacting with the TMD and therefore activation of the receptor^9^. Our work here presents an alternative view, in that they interact with the linker to induce a conformation within the GDxxGWxxxF motif that forms the tethered agonist. To test the role of these residues in stabilizing the LDLa-linker/relaxin complex, we prepared the double-mutant RXFP1_(1-72)_-L7K/K17A. ^15^N-CPMG relaxation dispersion experiments of RXFP1_(1-72)_-L7K/K17A in the apo-state show no residues within the LDLa have dispersion, but the linker residue Tyr53 retained dispersion with similar kinetics (k_ex_ 940 s^-1^, 2% minor state) to wild-type (Table 2). On titration of RXFP1_(1-72)_-L7K/K17A with relaxin-NH_2_, the majority of residues of the linker, with the notable exception of Trp46, that showed dispersion in wild-type also show dispersion with similar kinetics (k_ex_ 700 s^-1^, 3% minor state), however no residues within the LDLa show dispersion. As the derived apparent dissociation constants of the linker residues of both wild-type (25 μM) and RXFP1_(1-72)_-L7K/K17A (30 μM) are similar we conclude that this apparent K_D_ predominantly reflects both the stabilization of helical structure and the association with relaxin-NH_2_. Furthermore, the loss of all dispersion in the mutant LDLa supports that the dispersion observed in wild-type is indicative of stabilization and association of the linker, supporting that the major functional role of the LDLa is to stabilize structure within the linker for receptor activation.

Similar ^15^N-CPMG relaxation dispersion experiments were performed for RXFP2_(1-65)_ (Supplementary Fig. 5c). In the absence of relaxin-NH_2_, two residues show dispersion, Ser5 (k_ex_ 112 s^-1^, 8% minor state) in the disordered N-terminal of the LDLa and Gly42 (k_ex_ 71 s^-1^, 11% minor state) in the linker C-terminal to the LDLa module (Table 2). This dispersion likely reflects an interaction between these regions as previously described^20^. On addition of relaxin-NH_2_, these residues continue to show dispersion, but with the addition of His25 of the LDLa module and Trp47 of the linker. These residues show minor-state populations of 25-36% (Table 2), however, as all residues except Ser5 show chemical shift changes in the presence of relaxin-NH_2_ (Fig. 1), these data are consistent with the direct weak interaction of non-saturating relaxin-NH_2_ and not a conformational change.

## Discussion

Most class A GPCRs are activated by the binding of an agonist into an orthosteric site of the TMD shifting the conformational equilibrium to favour the active state. While this mechanism appears simple, variations of the nature of the agonist are extensive, especially for peptide- and protein-activated GPCRs. The LGR family, that includes the GHPRs and RXFP1/2, presents complex activation mechanisms that appear to utilize internal agonists, but each receptor shows significant differences. The current mechanism for the GPHRs^21^ proposes that binding of the hormone to the LRR domain induces a conformational change in a C-terminal hinge, structurally rearranging a 10-residue peptide, considered the agonist, to modulate the ECL region of the TMD leading to receptor activation. In contrast, for RXFP1 and RXFP2, binding of the hormone induces conformational changes in the N-terminal region of the receptor. Truncation of the N-terminal LDLa and analysis of N-terminal splice variants suggested that this module is the internal or tethered agonist for these receptors^14^. Mutagenesis experiments and receptor signalling assays suggested that key residues for RXFP1 were in the N-terminal region of the LDLa^8, 9^ while those for RXFP2 were in the C-terminal^10^, proposing that the role the LDLa played in activation differed for the two receptors. Importantly, despite extensive mutagenesis studies no essential residues of the LDLa were identified to implicate a direct role of the module in activation. However, mutation of the conserved residues within the GDxxGWxxxF motif, immediately C-terminal of the LDLa in both RXFP1^11^ and RXFP2^12^, did have significant effects on activation suggesting that these residues along with the LDLa may constitute the internal agonist. In our present study, we demonstrate that the function of the LDLa is not to directly activate the receptors, but to interact with the GDxxGWxxxF motif to modulate its shape and interaction with the TMD, and thus trigger receptor activation.

To understand how this internal agonist presents itself in the receptor we compared the structural relationship of the GDxxGWxxxF motif with the LDLa in the two receptors and used the common ligand, relaxin-NH_2_, to probe differences in binding sites and interactions. We propose that the motif in RXFP1 undergoes significant conformational change upon binding and activation by H2 relaxin (Fig. 7). The motif forms a loosely structured loop due to the association of the helical region, Leu48-Thr61, of the linker with the LDLa. Residues of the linker weakly interact with residues in the N-terminal region of the module, for example, Leu7 or Lys17. Mutagenesis of Leu7 and/or Lys17 lead to a weakening of both the linker/LDLa interaction and binding of H2 relaxin, and consequently a reduction in efficacy of activation as previously observed^9^. Therefore, in the context of the whole receptor, this helical region is likely to become well-structured upon H2 relaxin binding, due to interaction with both the LDLa and H2 relaxin. The GDxxGW portion of the motif does not directly participate in H2 relaxin binding, however, mutagenesis of the conserved residues, Asp42, Gly45 and Trp46, affect how the helical region of the linker associates with the LDLa, thus indirectly impacting on H2 relaxin affinity. Nevertheless, mutation of these residues results in a profound loss of activation, pointing to critical roles in TMD interactions. In contrast, in RXFP2 the GDxxGWxxxF motif is intimately associated and structured with the LDLa (Fig. 7). Its interaction with the module forms a binding-surface for H2 relaxin, involving residues in the C-terminal of the module, with little conformational change upon binding. For this receptor, mutation of the conserved residues, Asp43, Gly46 and Trp47 have a greater impact on H2 relaxin binding and activity than observed for RXFP1. However, these residues are likely to have similar key roles in receptor activation for both H2 relaxin and the cognate ligand of RXFP2, INSL3. Importantly, INSL3 does not interact with the linker or LDLa and the exact mechanism of activation for this ligand remains to be elucidated. Previous studies have highlighted that the N-terminus of the INSL3 A-chain is essential for RXFP2 activation^22^ suggesting that it may directly contact the extracellular loops or TMD of RXFP2 to enable the interaction of the GDxxGWxxxF motif for receptor activation^12^.

**Figure 7.**
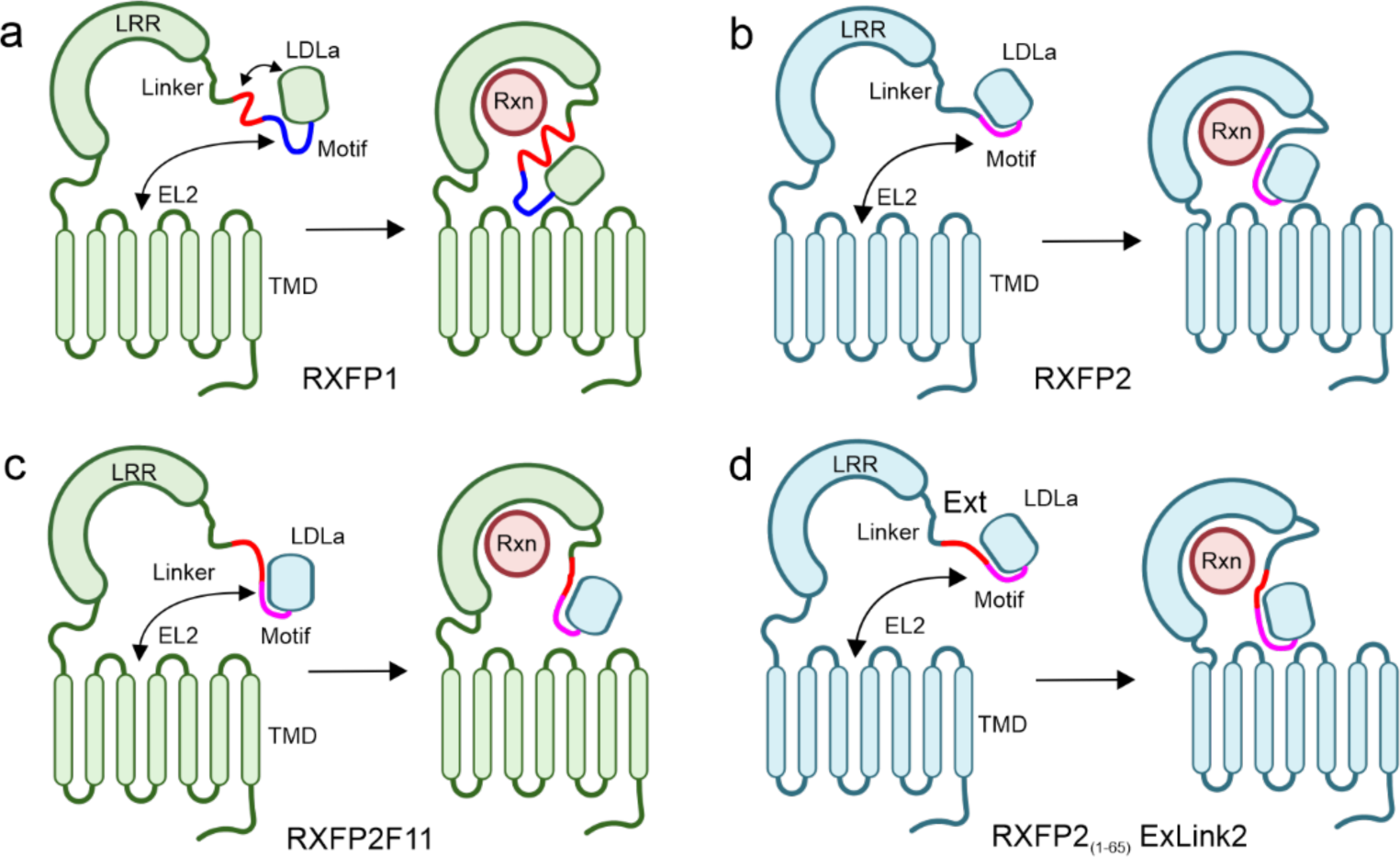
Schematic of the mechanisms of relaxin-mediated activation of RXFP1 and RXFP2. (a) **RXFP1**: In the absence of relaxin, the linker of RXFP1 weakly associates with the LDLa and this complex weakly interacts with EL2 of the transmembrane domain (TMD). The association of the linker with the LDLa induces a helical turn within the linker (red), while the GDxxGWxxxF motif (blue) remains largely unstructured. Relaxin associates strongly with the LRR domain, and weakly with the linker and not the LDLa. In combination the LDLa and relaxin stabilize and extend the linker helix (red), enabling the motif to form a conformation that leads to receptor activation. (b) **RXFP2**: In the absence of relaxin, the GDxxGWxxxF motif (magenta) closely associates with the LDLa. The LDLa-motif interacts with both EL2 of the TMD and relaxin. While little conformational change to the motif occurs, the strong association of relaxin with the LRR domain leads to the essential interaction of the LDLa-motif for receptor activation. (c) **RXFP2F11**: Replacing the LDLa-motif in RXFP1 with the RXFP2 LDLa-motif results in a receptor that weakens the binding of relaxin (20-fold) and reduces potency by 3000-fold. We propose that the structure of the RXFP2 LDLa-motif cannot effectively engage with relaxin or EL2 in this chimera (d) **RXFP2_(1-65)_ ExLink2**: Insertion of the 7-residues that encompass the relaxin binding site of RXFP1 into RXFP2 results in a chimera that behaves more like RXFP1, although helical structure is not induced in the linker. The chimeras highlight the different roles of the LDLa-motif in linker structure formation and stabilization in both receptors. Figure created with BioRender.com.

The difference in structure and interaction of the GDxxGWxxxF motif, as a requirement for H2 relaxin binding and activation, is reinforced by the behaviour of the construct RXFP2F11. Compared to the RXFP211 chimera, RXFP2F11 showed ∼3000-fold loss in H2 relaxin potency and >10-fold loss in H2 relaxin affinity. The only differences between these two chimeras are the non-conserved residues of the motif, which previously when mutated did not affect ligand-induced activation of RXFP2/H2 relaxin or RXFP2/INSL3^12^. Sequence alignments (Supplementary Fig. 6) show these non-conserved residues are conserved in the context of RXFP1 or RXFP2. Importantly, for organisms where no H2 relaxin gene has been identified but an INSL3 gene has, for example cow and rabbit, the residues are poorly conserved in RXFP1 but well conserved in RXFP2 pointing to the importance of these residues in maintaining the shape of the motif within the receptor. While very little is known about the structures of these sister receptors, our present study provides critical structural evidence of the differences in their H2 relaxin binding and activation mechanism. By studying the linker, a region that potentially differentiates these two receptors, we can bring to light some of the ways in which these two highly similar receptors are able to behave differently to one another in the biological context and explain how H2 relaxin has evolved into a cognate/specific ligand for RXFP1.

## Acknowledgements

This research was supported by National Health and Medical Research Council of Australia project grant 1100676 (R.A.D.B., D.J.S, M.D.W.G and P.R.G.), the Victorian Government Operational Infrastructure Support Program and equipment grants from the Australian Research Council (LE120100022). R.A.D.B. is supported by an NHMRC Research Fellowship.

## Methods

### Peptides

Native recombinant H2 relaxin (serelaxin) was kindly provided by Corthera Inc. Amidated H2 relaxin with both the A and B-chain C-termini amidated, was synthesized using Fmoc chemistry and regioselective disulfide bond formation as previously described for amidated H2 relaxin and other analogues^13, 23^. After synthesis, the peptide was purified and characterized by reverse phase RP-HPLC and MALDI-TOF MS.

### Protein Expression and Purification

The DNA sequences of RXFP1_(1-72)_, RXFP2_(1-65)_, RXFP1_(1-40)_ and RXFP2_(1-41)_ were amplified by PCR and inserted into the vector pGEV2, an N-terminal thrombin-cleavable GB1 fusion expression vector using BamHI and XhoI restriction sites. For RXFP2_(1-65)_, Pro4 was mutated to phenylalanine to remove cis-trans isomerism as previously reported^12^. All site-specific mutants were prepared using PrimeStar DNA Taq Polymerase (Takara Clonetech). Expression and purification of recombinant wild-type and mutant RXFP1_(1-72)_, RXFP2_(1-65)_, RXFP1_(1-40)_ and RXFP2_(1-41)_ followed our previous methods with minor modifications^8–11^. Briefly, proteins were expressed into BL21(DE3) trxB (Novagen) using autoinduction^24^, for producing unlabelled protein samples for SAXS experiments. For uniform ^15^N isotopic labelling, protein was expressed in N5052 minimal medium^24^ using ^15^NH_4_Cl (Sigma-Aldrich). For ^13^C,^15^N labelling, cells were grown in a 1-L Braun Biostat fermenter supplemented with ^15^NH_4_Cl and D-[^13^C] glucose (Sigma-Aldrich) as the sole source of nitrogen and carbon for the bacterial cells, as suggested by Cai et al.^25^ Cells were harvested, pelleted, and stored at −20 °C. Following resuspension in 50 mM Tris, 150 mM NaCl, 5mM EDTA-Na and 1mM phenylmethylsulfonyl fluoride (PMSF) at pH 7.4, cell pellets were lysed using an Avestin EmulsiFlex C3 cell crusher. Cell debris was removed by centrifugation (13,000 × g, 4 °C for 40 min), the soluble fraction was filtered using a 0.45 µm syringe filter. The GB1-LDLa linker fusion proteins were purified over IgG Sepharose 6 Fast Flow beads, eluted with 50mM acetic acid, pH 3.4. Further, the eluted protein was buffer exchanged (50 mM Tris-HCl, 150mM NaCl, pH 8.5) via dialysis, and refolded (100–300 mg.ml^-1^, 3 mM GSH, 0.3 mM GSSG, 50mM Tris-HCl, 150 mM NaCl, 5 mM CaCl_2_, pH 8.5) by incubating overnight at 4 °C with gentle stirring. The GB1 tag was cleaved from the oxidized proteins by incubating overnight with thrombin (10 U.mg^−1^ protein, Sigma-Aldrich). The cleaved protein was purified by reversed phase high-performance liquid chromatography (RP-HPLC) (buffer A: 0.1% trifluoro-acetic acid in water, buffer B 100% acetonitrile with 0.1% trifluoro-acetic acid; Agilent Zorbax 300SB-C18 column). Collected protein fractions were lyophilized and stored at −20 °C.

The expression and purification of RXFP2_(1-65)_ ExLink 2 was performed as previously described^17^. Protein was expressed as a His_6_-tag fusion in BL21(DE3) trxB (Novagen) using autoinduction^24^. ^15^N and ^13^C,^15^N labelling were performed as described above. Cells were harvested, lysed and purified using Talon Superflow resin (Takara Clontech). Further, the eluted protein was refolded using the reduced and oxidized glutathione and His_6_-GB1 tag was cleaved using thrombin, followed by RP-HLC, also described above.

The peptide mimetics of the exoloops, ssRXFP1 (EL1^(475-486)^/EL2-RXFP1) and ssRXFP2 (EL1^(475-486)^/EL2-RXFP2), were purified as previously described^11, 15^. Proteins were expressed from BL21(DE3) as His_6_-tag fusions. Expression was induced by isopropyl –D-1-thiogalactopyranoside (IPTG) induction in LB medium for 16 h at 16 °C. Cells were harvested, pelleted, lysed and purified over Talon Superflow resin (Takara Clontech). The His_6_-tag was removed by thrombin cleavage and the proteins further purified with a HiLoad 16/60 Superdex 75 prep grade column (GE Healthcare) in 20 mM Tris HCl (pH 7.4), 150 mM NaCl.

### NMR spectroscopy

The ^15^N-labeled RXFP1_(1-72)_ sample, which comprises the first 72 residues of the receptor, was identical to the material used for the assignment of the backbone chemical shifts as previously described^26^. For ^15^N RXFP2_(1-65)_ and RXFP2_(1-65)_ ExLink2, the assignments were adapted from previously reported assignments for C-terminal GB1 fused RXFP2_(1-65)_ and RXFP2_(1-65)_ ExLink2 construct^12, 17^. All protein samples were prepared in a buffer containing 50 mM Imidazole, 10 mM CaCl_2_ at pH 6.8 for all experiments. Protein samples used for titrations were dialyzed in the same buffer and same vessel overnight to ensure sample buffer conditions. NMR spectra were acquired on a Bruker Avance II 800 MHz spectrometer equipped with TXI cryoprobe and a single axis field gradient (Gz). NMR spectra were processed using NMRPipe ^27^ as described in ^26^ and data analyzed in NMRFAM-SPARKY ^28^. The ^1^H chemical shifts were referenced directly to DSS at 0 ppm and the ^13^C and ^15^N chemical shifts were subsequently referenced using the ^13^C/^1^H and ^15^N/^1^H ratios ^29^. The ΔCα – ΔβC smoothed values were calculated as previously^11, 30^. The ^15^N,^1^H chemical shift mapping was computed using ^31^:

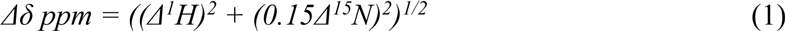

### SAXS data collection and analysis

Small-angle X-ray scattering (SAXS) measurements were conducted at the Australian Synchrotron SAXS/WAXS Beamline equipped with a co-flow system to avoid radiation damage and enable higher X-ray flux (11,500 eV) and with optimized in-line size exclusion chromatography (SEC) to limit protein sample dilution ^32–34^. Fifty microliters of the purified RXFP1_(1-72)_ and RXFP2_(1-65)_ at 5 mg/ml was injected over a Superose 6 5/150 increase column (GE Healthcare) equilibrated and eluted with TBS (pH 7.4) with a solution containing 10 mM calcium chloride and 0.2% sodium azide. The sample to detector length used was 1426 mm, providing a q range of 0.007 - 0.515 Å-1. Collected SAXS data was reduced using the Sactterbrain software, analysed by CHROMIXS^35^, and the ATSAS software package 3.0.2 (r12592)^36^. SAXS patterns, the radius of gyration (*R*_g_), the maximal particle dimension (*D*_max_), and the pairwise distance distribution histogram [P(*r*) plot] were determined by using the ATSAS software^36^. *Ab initio* modelling was performed using DAMMIN^37^. Ensemble of conformations and models were generated using EOM 2.0^19^ in the ATSAS software suite. A summary of the SAXS data acquisition parameters is provided in Supplementary Table 2.

### Construction of RXFP2F11

The chimeric construct RXFP2F11 consists of the RXFP2 sequence up to and including Phe51, followed by the RXFP1 sequence from Asp51 to the end of the TM domain. It was made using overlap PCR by the methods described in ^20^, with primers Fwd: CATCATGGATCCGCCACCATGGACAGCAAAG and Rev: 5’ CATTTTGTAGTAACTGGCAAAATATTTGTCAAATATGGTCGCCCATCCACTAGTG 3’ on RXFP2 template, and Fwd: 5’ CACTAGTGGATGGGCGACCATATTTGACAAATATTTTGCCAGTTACTACAAAATG 3’ and Rev: GAGAGCTCGAGTCATGAATAGGAATTGAGTCTCGTTG on RXFP1 template to make the two DNA portions. The final annealed product was cut with BamHI and XhoI restriction enzymes for insertion into a freshly cut pcDNA3.1TM/Zeo+ AmpR mammalian expression vector (Invitrogen, Carlsbad, CA, USA) and then the entire insert sequenced to ensure accuracy.

### Receptor expression in HEK293T cells

HEK293T cells (ATCC #CRL-1573; American Type Tissue Culture Collection) were used for receptor expression and were grown in Dulbecco’s modified eagle medium supplemented with 10% foetal bovine serum, 1% l-glutamine and 1% penicillin/streptomycin in incubators maintained at 37 °C with 5% CO_2_ and 85% humidity. Transient transfections were achieved using lipofectAMINE 2000 (Invitrogen) according to the manufacturer’s instructions.

### Binding and signaling assays

The affinity of relaxin for the mutant receptors in comparison to wild-type RXFP1 was assessed using Europium (Eu^3+^)-labelled H2 relaxin (Eu-H2) ^38^ at increasing concentrations in the presence or absence of 1 μM unlabeled ligand. Following a 1 h incubation, media was removed and 100 μl Delfia Enhancement solution (PerkinElmer) was added to each well. Time-resolved fluorescence with excitation at 340 nm and emission at 614 nm was read on an Omega POLARstar plate reader after incubation in low light for 20–30 mins with shaking. Data from at least three independent experiments, all performed in triplicate were pooled and presented as mean fluorescent specific binding ± SEM. Binding data was analysed by nonlinear regression one-site binding curves using GraphPad PRISM 6 to calculate K_D_ values. cAMP activation in response to ligand stimulation was measured by co-transfection with a pCRE β-galactosidase reporter gene as previously described ^14^. Briefly, co-transfected cells were stimulated with increasing concentrations of H2 relaxin and 5 μM Forskolin or media only were used as a positive and negative controls, respectively. Following a 6 h incubation at 37 °C media was aspirated and plates frozen at −80 °C for ≥ 24 h. β-galactosidase activity in plates was achieved as previously described ^14^ with absorbance readings measured on a Benchmark Plus Microplate Reader (Bio-Rad) at 570 nm. All experiments were performed in triplicate a minimum of three times and data were pooled and presented as percentages of the 5% Forskolin response. GraphPad PRISM was used to fit a nonlinear regression sigmoidal dose-response curve and resulting pEC50 and maximum response (Emax) values were subjected to one-way ANOVA and uncorrected Fisher’s least square difference comparison test.

### Cell surface expression assays

The presence of mutant receptors at the surface of cells was gauged by virtue of the FLAG epitope present on their N-termini using a plate-based ELISA assay. HEK293T cells were seeded at 2 × 10^5^ cells per well in 24-well plates (Costar) pre-coated with poly(l-lysine) and transfected with 1 mg per well of plasmid DNA. After a further 24 h of growth, cells were washed in TBS/CaCl_2_ (50 mM Tris pH 7.4, 150 mM NaCl and 1 mM CaCl_2_) and fixed with 3.7% formaldehyde in TBS/CaCl_2_ for 20 min. Two washes followed and then incubation for 45 min at room temperature with 1% bovine serum albumin (BSA) in TBS/CaCl2 to block non-specific binding. The cells were then incubated with 10 mg.ml^-1^ of anti-FLAG M1 monoclonal antibody (Sigma-Aldrich) in TBS/CaCl_2_ for 2 h at room temperature. Cells were washed twice more before a 15-min reblock in 1% BSA/TBS/CaCl_2_; then incubated for 1 h with 2 mg.ml^-1^ goat anti-mouse Alexa Fluor 488 suspended in 1% BSA/TBS/CaCl_2_. Cells were then washed thrice and stored frozen at −80 °C overnight. Finally, cells were thawed and lysed by incubating for 30 min with 200 ml per well of lysis buffer (50 mM Tris pH 7.4, 150 mM NaCl, 1 mM EDTA, 0.25% Triton X-100) at room temperature with shaking and scraped and transferred to black 96-well plates (180 ml in each well; Costar) to be read on an Omega POLARstar plate reader (BMG labtech) with excitation at 490 nm and emission at 520 nm. Non-specific background was determined using cells transfected with empty vector and mutant receptor expression was expressed as the percentage of the wild-type receptor expression. Data are mean ± s.e.m. from at least three independent experiments each performed in triplicate. Pooled data were analysed in GraphPad PRISM 6 using one-way analysis of variance and uncorrected Fisher’s least square difference multiple comparison test.

### Backbone ^15^N relaxation parameters and reduced spectral density mapping

^15^N relaxation parameters R_1_, R_2_, and steady state ^15^N{^1^H}-NOE of RXFP1_(1-72)_ were measured at ^15^N frequency of 81.1 MHz with 1024 and 150 complex data points over spectral widths of 13.0 and 26.0 ppm for ^1^H (F2) and ^15^N (F1), respectively at pH 6.8 and at 15 and 25 °C, using pulse sequences with sensitivity enhancement provided in the Bruker pulse sequence library, as described previously^39–41^. R_1_ and R_2_ experiments were collected with a recycle time of 2.6 s and 16 scans per FID and ^15^N{^1^H}-NOE experiments were collected with a saturation pulse of 4 s and an additional relaxation delay of 5 s and 32 scans per FID. Relaxation delays of 10 (×2), 50, 100, 300 (×2), 500, 700, 900 (×2) and 1500 ms for R_1_ experiments and for R_2_ relaxation delays of 16.96 (×2), 33.92, 67.84, 101.76 (×2), 135.68, 169.6 (×2), 203.52 and 237.44 ms were used. The repeated spectra were used to estimate instrumental error. ^15^N relaxation parameters were determined using the program *relax* ^42–44^ (version 3.3.4). For R_1_ and R_2_ rate constants, errors were estimated using 500 Monte Carlo Simulations. The steady state ^15^N{^1^H}-NOE values for RXFP1_(1-72)_ were estimated from the ratios of peak intensities obtained from spectra acquired with and without proton saturation using *relax*. Errors for ^15^N{^1^H}-NOE experiment were calculated based on the noise level in the spectra.

### Reduced Spectral Density Mapping

The relaxation data acquired at ^15^N frequency of 81.1 MHz on the 800 MHz spectrometer was also analyzed using spectral densities ^41^. The reduced spectral density parameters were calculated using *Mathematica* ^45^; based on the following equations:

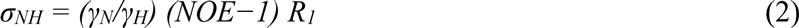

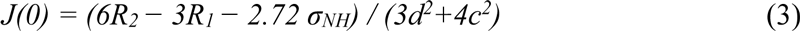

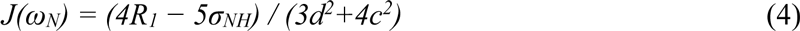

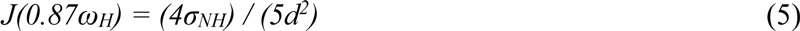

where J(0), J(ω_N_) and J(0.87ω_H_) are values of spectral density sampling frequencies ω = 0, ω_N_ (81 MHz), and ω_H_ (800 MHz), respectively. The other parameters are given by d = (μ_0_*h*γ_N_γ_H_/(8π^2^))/ < r^3^_NH_ >, c = ω_N_(Δσ)/√3, μ_0_ is the permeability of the free space, *h* is Planck’s constant, γ_H_ and γ_N_ are the gyromagnetic ratio of ^1^H and ^15^N respectively, r_NH_ is the average amide bond length (1.02 Å), and Δσ is the chemical shift anisotropy for ^15^N nuclei (−170 ppm).

### ^15^N-relaxation dispersion on RXFP1_(1-72)_

Backbone amide ^15^N Carr-Purcell-Meiboom-Gill (CPMG) constant time relaxation dispersion experiments were acquired on apo ^15^N labeled RXFP1_(1-72)_ and in complex with H2 relaxin at 18.8 and 16.5 T with 1024 and 128 complex data points over spectral widths of 13.0 and 26.0 ppm for ^1^H (F2) and ^15^N (F1), respectively at pH 6.8 and at 298 and 288 K. A series of interleaved two-dimensional spectra were collected at different υ_CPMG_ frequencies of 50, 75 (×2), 100, 150 (×2), 200, 300, 500, 700 (×2), 800, 900, 1000, 1500 MHz, with a total relaxation delay τ_CPMG_ of 80 ms. All NMR spectra were processed using NMRPipe ^27^ and analyzed in NMRFAM-SPARKY ^28^.

The Effective relaxation rates, R*_2,eff_*, were extracted by fitting the dispersion data to equation (6) using NESSY ^46^:

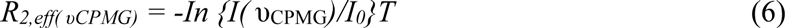

where *I(* υ_CPMG_) is the peak intensity in the spectrum recorded at various υ_CPMG_ values and *I_0_* is the peak intensity in the reference spectrum with no relaxation delay (τ_CPMG_ = 0 ms). *R_2,eff(υCPMG)_* values obtained for RXFP1_(1-72)_ in apo- and relaxin bound form at 15 °C were fitted to Carver-Richard equations for two-site exchange model undergoing slow exchange (*k*_ex_ << Δω) regime on the Sherekhan webserver ^47^.

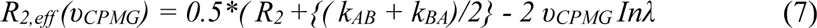

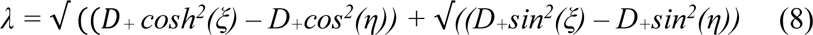

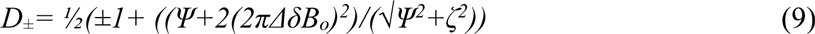

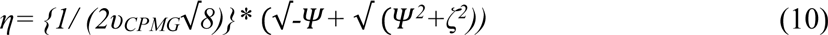

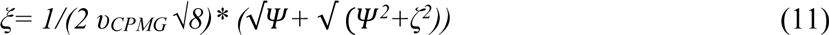

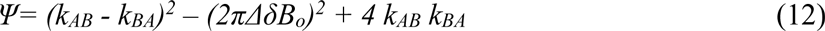

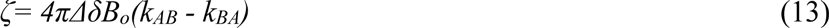

