## Supplementary figures for "Structural insights into the modes of relaxin-binding and tethered-agonist activation of RXFP1 and RXFP2"

**Supplementary Table 1**: $^{15}N$ dynamics parameters of apo- and relaxin-bound $RXFP1_{(1-72)}$ and $RXFP2_{(1-65)}$

| $RXFP1_{(1-72)}$ | H2 | Temp ºC | $^{15}N\{^1H\}$ NOE | $^{15}N$-$R_1$ $s^{-1}$ | $^{15}N$-$R_2$ $s^{-1}$ | J(0) ns.rad$^{-1}$ | $J(0.87\ \omega_H)$ ps.rad$^{-1}$ | $J(\omega_N)$ ns.rad$^{-1}$ |
|---|---|---|---|---|---|---|---|---|
| LDLa (6 to 40) | - | 25 | 0.78 ± 0.08 | 1.4 ± 0.1 | 6.7 ± 1.1 | 2.3 ± 0.3 | 6.1 ± 1.4 | 0.34 ± 0.03 |
| Linker (41 to 60) | - | 25 | 0.53 ± 0.09 | 1.4 ± 0.1 | 4.9 ± 1.4 | 1.5 ± 0.4 | 10.6 ± 1.2 | 0.33 ± 0.03 |
| LDLa (6 to 40) | + | 25 | 0.78 ± 0.08 | 1.4 ± 0.2 | 7.3 ± 1.2 | 2.5 ± 0.4 | 5.7 ± 1.4 | 0.33 ± 0.04 |
| Linker (41 to 60) | + | 25 | 0.63 ± 0.10 | 1.1 ± 0.2 | 8.6 ± 2.8 | 3.1 ± 1.1 | 7.1 ± 2.5 | 0.27 ± 0.05 |
| LDLa (6 to 40) | - | 15 | 0.83 ± 0.06 | 1.1 ± 0.1 | 10.0 ± 2.4 | 3.7 ± 1.0 | 3.7 ± 1.5 | 0.27 ± 0.04 |
| Linker (41 to 60) | - | 15 | 0.59 ± 0.07 | 1.1 ± 0.2 | 9.4 ± 2.6 | 3.3 ± 0.9 | 6.9 ± 1.7 | 0.25 ± 0.05 |
| LDLa (6 to 40) | + | 15 | 0.84 ± 0.06 | 1.0 ± 0.1 | 11.4 ± 2.4 | 4.1 ± 0.8 | 2.8 ± 0.7 | 0.25 ± 0.03 |
| Linker (41 to 60) | + | 15 | 0.78 ± 0.07 | 0.6 ± 0.1 | 11.2 ± 2.6 | 4.1 ± 1.0 | 3.9 ± 1.4 | 0.21 ± 0.03 |
| $RXFP2_{(1-65)}$ | | | | | | | | |
| LDLa (5 to 41) | - | 25 | 0.76 ± 0.10 | 1.5 ± 0.1 | 7.8 ± 0.9 | 3.0 ± 0.4 | 5.4 ± 2.4 | 0.38 ± 0.04 |
| Linker (42 to 52) | - | 25 | 0.62 ± 0.10 | 1.5 ± 0.1 | 6.9 ± 1.2 | 2.5 ± 0.5 | 7.7 ± 1.8 | 0.33 ± 0.03 |
| LDLa (5 to 41) | + | 25 | 0.80 ± 0.09 | 1.4 ± 0.1 | 8.7 ± 2.0 | 3.3 ± 0.7 | 4.6 ± 2.5 | 0.35 ± 0.04 |
| Linker (42 to 52) | + | 25 | 0.69 ± 0.09 | 1.4 ± 0.1 | 9.1 ± 1.7 | 3.2 ± 0.8 | 6.4 ± 2.1 | 0.30 ± 0.04 |
| LDLa (5 to 41) | - | 15 | 0.81 ± 0.08 | 1.4 ± 0.1 | 8.4 ± 0.8 | 3.3 ± 0.4 | 4.7 ± 2.3 | 0.36 ± 0.06 |
| Linker (42 to 52) | - | 15 | 0.74 ± 0.08 | 1.4 ± 0.0 | 8.6 ± 1.7 | 3.4 ± 0.6 | 6.0 ± 1.8 | 0.29 ± 0.03 |
| LDLa (5 to 41) | + | 15 | 0.83 ± 0.07 | 1.3 ± 0.1 | 9.9 ± 1.8 | 4.0 ± 0.6 | 3.3 ± 2.2 | 0.32 ± 0.05 |
| Linker (42 to 52) | + | 15 | 0.78 ± 0.05 | 1.3 ± 0.1 | 10.4 ± 1.5 | 4.1 ± 0.6 | 5.0 ± 1.8 | 0.28 ± 0.04 |

**Supplementary Table 2**: SAXS results for $RXFP1_{(1\text{-}72)}$ and $RXFP2_{(1\text{-}65)}$.

| **SAXS data collection parameters:** | **$RXFP1_{(1\text{-}72)}$** | **$RXFP2_{(1\text{-}65)}$** |
|---|---|---|
| Instrument/source | Australian Synchrotron SAXS/WAXS beamline equipped with Pilatus 1M detector and sheath-flow cell for SEC-SAXS. | |
| Wavelength (Å) | 1.0332 | |
| Beam energy (kEv) | 12 | |
| Beam size (µm) | 250 × 130 | |
| Sample-to-detector distance (mm) | 1426 | |
| $q$ ($Å^{-1}$) | 0.005 – 0.334 | |
| Absolute scaling method | Comparison with scattering from 1 mm pure water | |
| Normalization | To transmitted intensity from beamstop counter | |
| Exposure time | 1 s measurements from SEC-SAXS elution | |
| Sample temperature (K) | 295 | |
| **SEC-SAXS parameters** | | |
| Column | Superdex 200 5×150 | |
| Flow rate (mL/min) | 0.4 | |
| Concentration (mg/mL) | 5 | |
| Injection volume (µL) | 50 | |
| Average conc. (mg/mL) | 3.02 | |
| Solvent | 20 mM Tris-HCl, pH 7.4, 150 mM NaCl, 10 mM $CaCl_2$, 0.1% $NaN_3$ | |
| **Software employed** | | |
| SAXS data reduction | $I(q)$ vs $q$ using Scatterbrain 3.0.2, SEC-SAXS solvent subtraction using *CHROMIXS* from *ATSAS 3.0.2* | |
| Basic analysis (Guinier, *P(r)*, molecular mass) | *PRIMUSqt* from *ATSAS 3.0.2* | |
| *Ab initio* modelling | *DAMMIN & DAMMIF* from *ATSAS 3.0.2* | |
| Atomic structure modelling | *EOM 2.0* from *ATSAS 3.0.2* | |
| **Structural parameters** | | |
| Mass from $V_c$ (kDa) (expected mass, ratio to expected in brackets) | 7.68 (8.29, 0.93) | 6.35 (7.09, 0.89) |
| **Guinier analysis** | | |
| $R_g$ (Å) | 20.74 ± 0.19 | 17.12 ± 0.14 |
| *I(0)* ($cm^{-1}$) | 0.48 ± 0.00025 | 0.41 ± 0.00038 |
| $qR_g$ min,max | 0.27, 1.24 | 0.21, 1.29 |
| ***P(r)* analysis** | | |
| $R_g$ (Å) | 21.84 ± 0.19 | 18.12 ± 0.23 |
| *I(0)* ($cm^{-1}$) | 0.4876 ± 0.000243 | 0.4144 ± 0.000103 |
| $D_{max}$ (Å) | 78.51 | 71.04 |
| Porod volume ($Å^3$) | 19957.10 | 12151.60 |
| **Atomic modelling** | | |
| ***EOM*** | | |
| Input structure | NMR structure for RXFP1 (2jm4) and RXFP2 (2m96) | |
| Symmetry assumptions | P1 (LDLa domain and flexible residues) | |
| Flexible residues modelled | 32 residues at C terminus (RXFP1 linker), 25 residues at C-terminus (RXFP2 linker) | |
| Flexible residues | 32 residues C-terminus | 25 residues C-terminus |
| $\chi^2$ range | *1.2-1.3* | *0.98-1.1* |

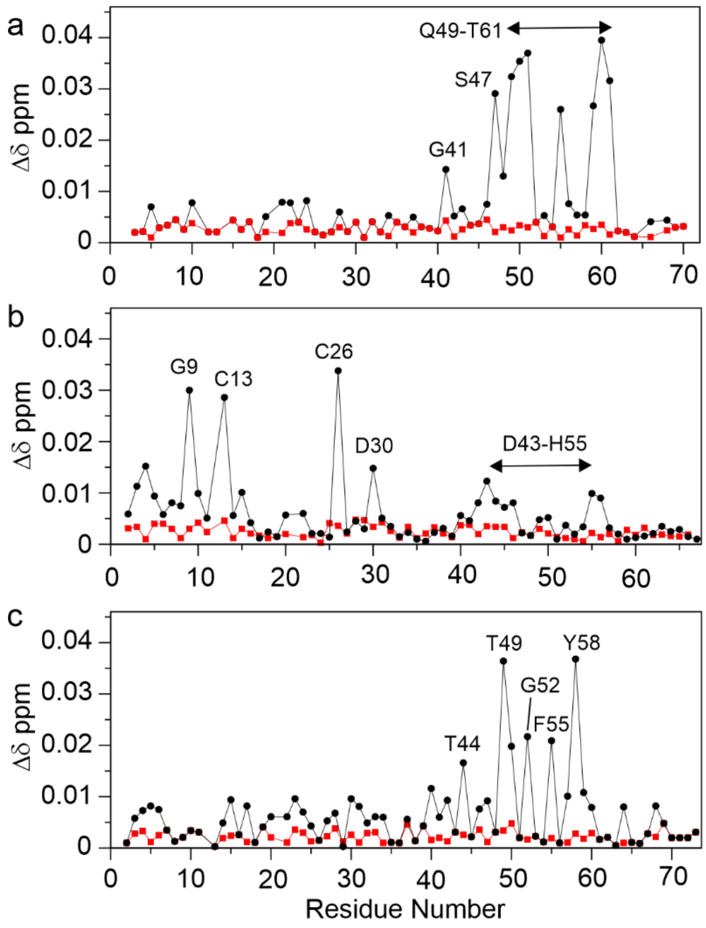


**Supplementary Figure 1: Interactions of RXFP1$_{(1\text{-}72)}$, RXFP2$_{(1\text{-}65)}$ and RXFP2$_{(1\text{-}65)}$ ExLink2 with Transmembrane Domain exoloop mimetics.** Titration of (a) 25 μM $^{15}$N-labelled RXFP1$_{(1–72)}$ with twenty molar equivalents of EL1$^{(475\text{-}486)}$/EL2-RXFP1in the presence (black circles) and absence (red squares) of 10 mM $CaCl_2$. (b) 25 μM $^{15}$N-labelled RXFP2$_{(1–65)}$ with twenty molar equivalents of EL1$^{(475\text{-}486)}$/EL2-RXFP2 in the presence (black circles) and absence (red squares) of 10 mM $CaCl_2$. (c) 25 μM $^{15}$N-labelled RXFP2$_{(1–65)}$ ExLink2 with twenty molar equivalents of EL1$^{(475\text{-}486)}$/EL2-RXFP2 in the presence (black circles) and absence (red squares) of 10 mM $CaCl_2$. Experiments were conducted at pH 6.8 and 25 ºC.

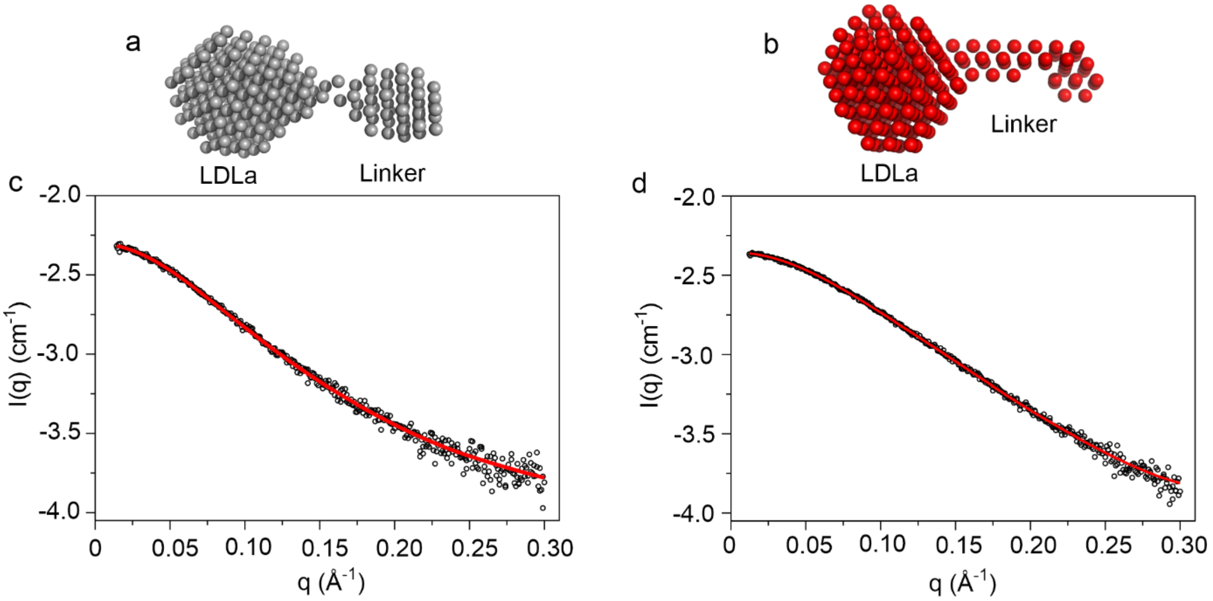


**Supplementary Figure 2: An initio shape modelling of RXFP1$_{(1-72)}$ and RXFP2$_{(1-65)}$.** Representative single-bead models from DAMMIN are shown in beads of (a) RXFP1$_{(1-72)}$ and (b) RXFP2$_{(1-65)}$. These data support a globular RXFP1$_{(1-72)}$ and extended RXFP2$_{(1-65)}$. Plots of the DAMMIN model against the experimental scattering curve for (c) RXFP1$_{(1-72)}$ and (d) RXFP2$_{(1-65)}$.

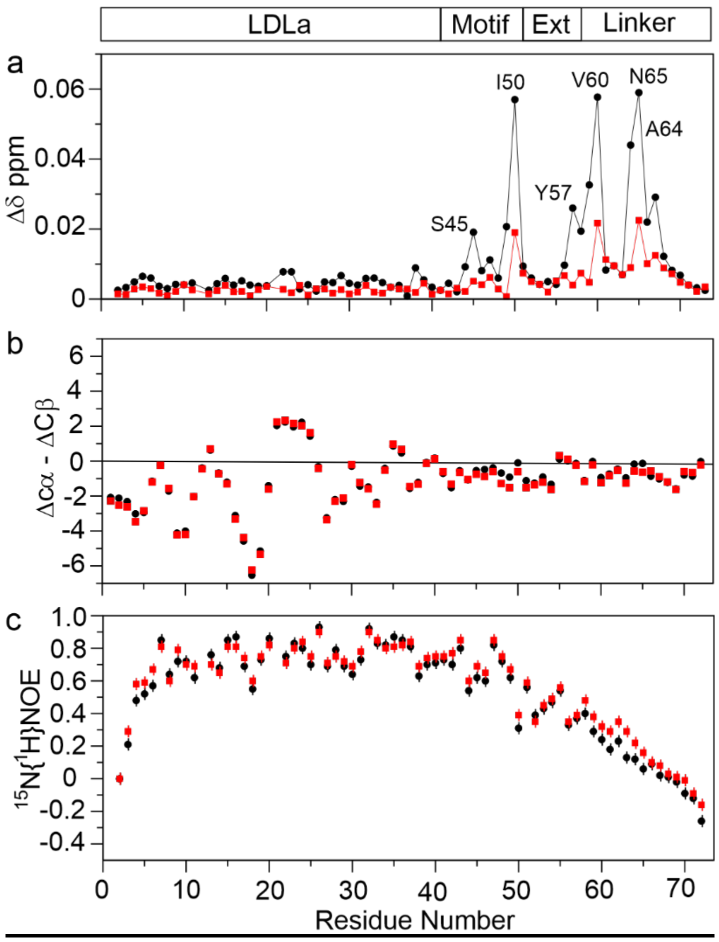


**Supplementary Figure 3**: **NMR analysis of RXFP2$_{(1-65)}$ with the 7-residue relaxin binding site of RXFP1 inserted (RXFP2$_{(1–65)}$ ExLink2).** (a) Titration of 25 µM $^{15}$N-labelled RXFP2$_{(1–65)}$ ExLink2 with twenty equivalents of EL1$^{(475-486)}$/EL2-RXFP2 in the presence (black circles) and absence (red squares) of 10 mM $CaCl_2$; (b) Plots of $^{13}$Cαβ secondary chemical shifts; (c)$^{15}$N{$^{1}$H}-NOEs in apo- (black circle) and in the presence of 20 equivalents of relaxin (red square) for RXFP2$_{(1–65)}$ ExLink2. Experiments were conducted at pH 6.8 and 25 ºC. The sequence location of the LDLa module, GDxxGWxxxF motif, the 7-residue insertion (Ext, Lys52-Tyr58) and the remainder of the linker are indicated.

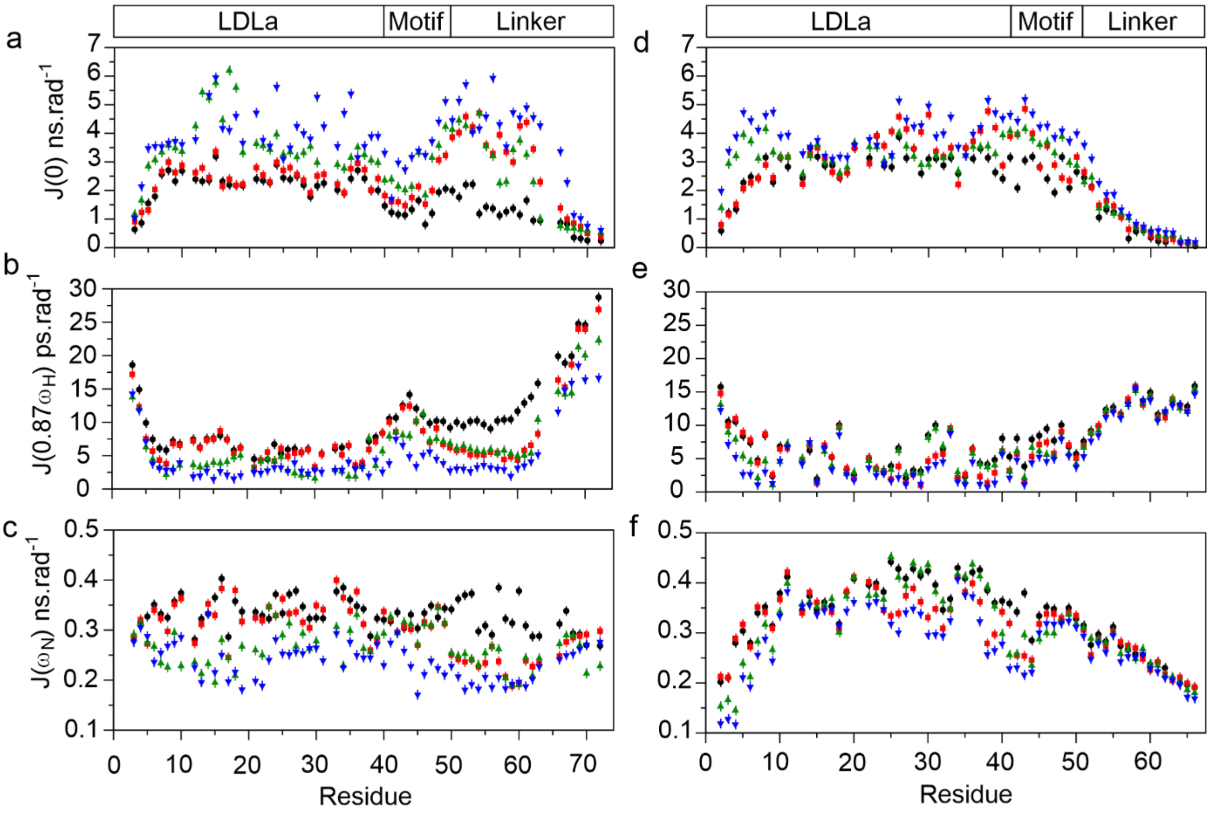


**Supplementary Figure 4: Reduced spectral density analysis of RXFP1$_{(1\text{-}72)}$ and RXFP2$_{(1\text{-}65)}$.** Analysis of 100 µM RXFP1$_{(1\text{-}72)}$ (a-c) and RXFP2 $_{(1\text{-}65)}$ (d-f) for the apo-state at 25 ºC (black circles), and at 15 ºC (green triangle), with amidated relaxin at 25 ºC (red square) and at 15 ºC (blue triangle) measured at $^{15}$N frequency of 81.1 MHz and at pH 6.8. Sample concentrations represent 83% saturation of RXFP1$_{(1\text{-}72)}$ and RXFP2$_{(1\text{-}65)}$ with ligand. The sequence locations of the LDLa module, the GDxxGWxxxF motif and the remainder of the linker in the receptors are shown above (a) and (d).

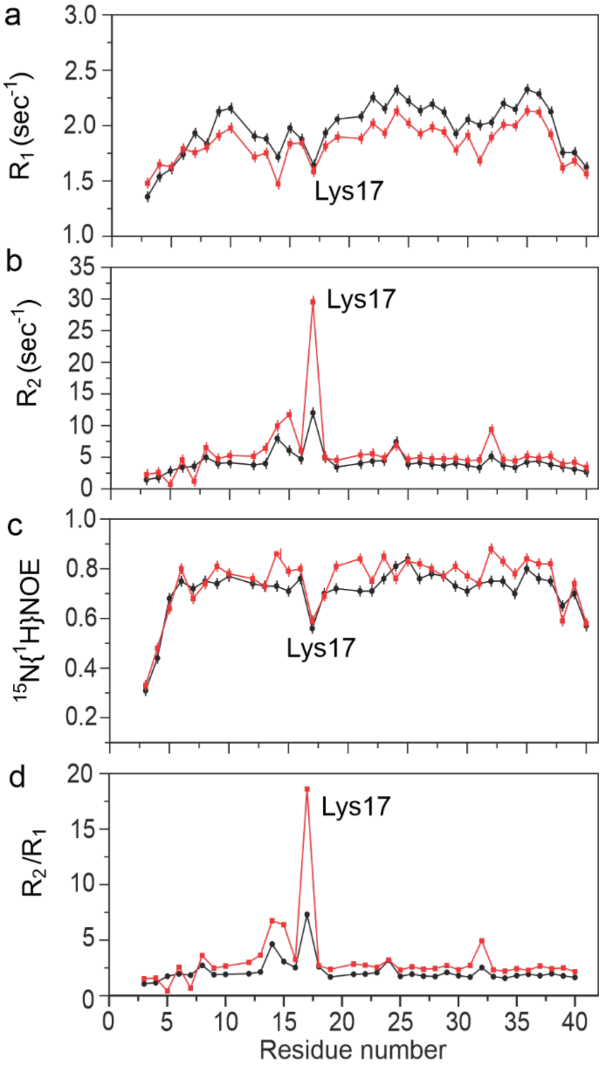


**Supplementary Figure 5**: **$^{15}$N-spin relaxation dynamics of the RXFP1 LDLa module.** Backbone $^{15}$N relaxation parameters of 100 µM RXFP1$_{(1\text{-}40)}$ at 25 ºC (dark circles) and at 15 ºC (red square) measured at $^{15}$N frequency of 81.1 MHz and pH 6.8. (a) $^{15}N$ $R_1$ (b) $^{15}N$ $R_2$ (c) Steady State $^{15}N\{^1H\}$ NOE (d) $R_2/R_1$ ratio. Error bars are calculated using Monte Carlo Simulations for $R_1$ and $R_2$ measurements and based on average estimated noise level for $^{15}N\{^1H\}$ NOE.

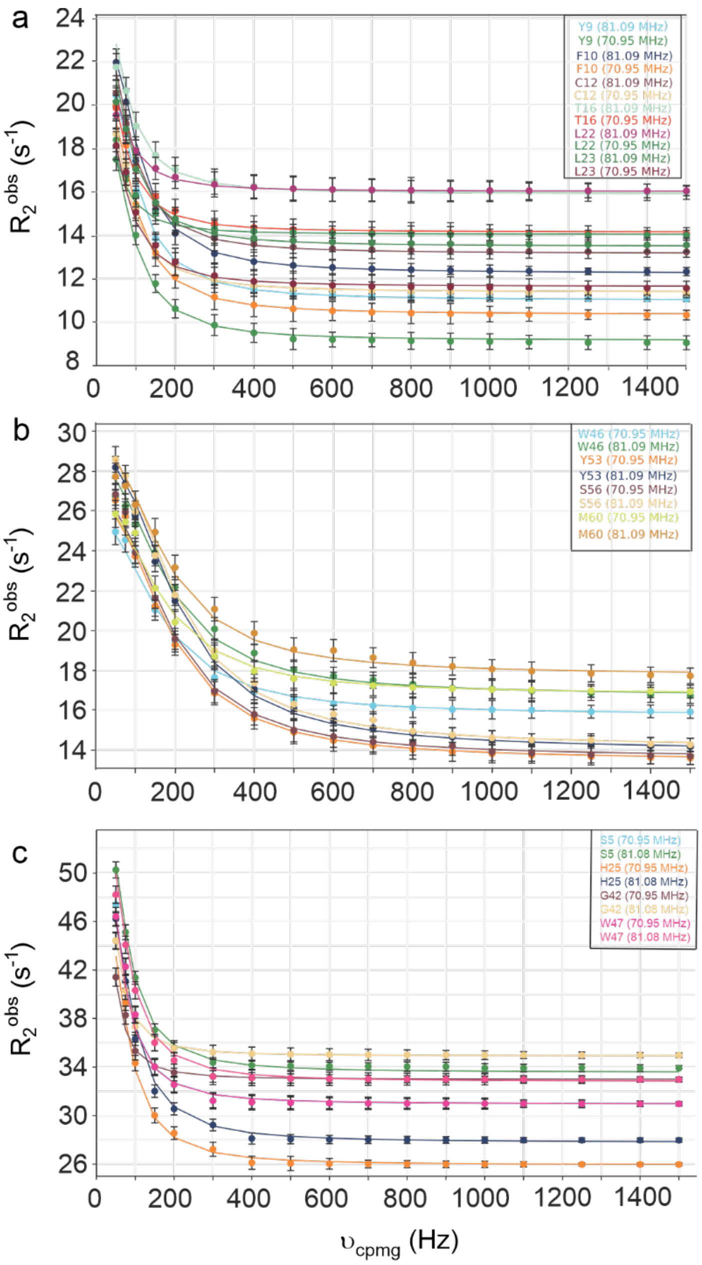


**Supplementary Figure 6**: **$^{15}$N Relaxation dispersion analysis of RXFP1$_{(1-72)}$ and RXFP2$_{(1-65)}$.** $^{15}$N CPMG relaxation dispersion data of backbone amides of RXFP1$_{(1-72)}$ (a, b) and RXFP2 $_{(1-65)}$ (c) in the presence of relaxin at 15 ºC. Representative curves for residues of the RXFP1 (a) LDLa module and (b) linker region and (c) RXFP2 LDLa and linker region. $k_{ex}$ ($s^{-1}$), pB%, $k_{on}$ and $k_{off}$ rates are provided in Table 3.

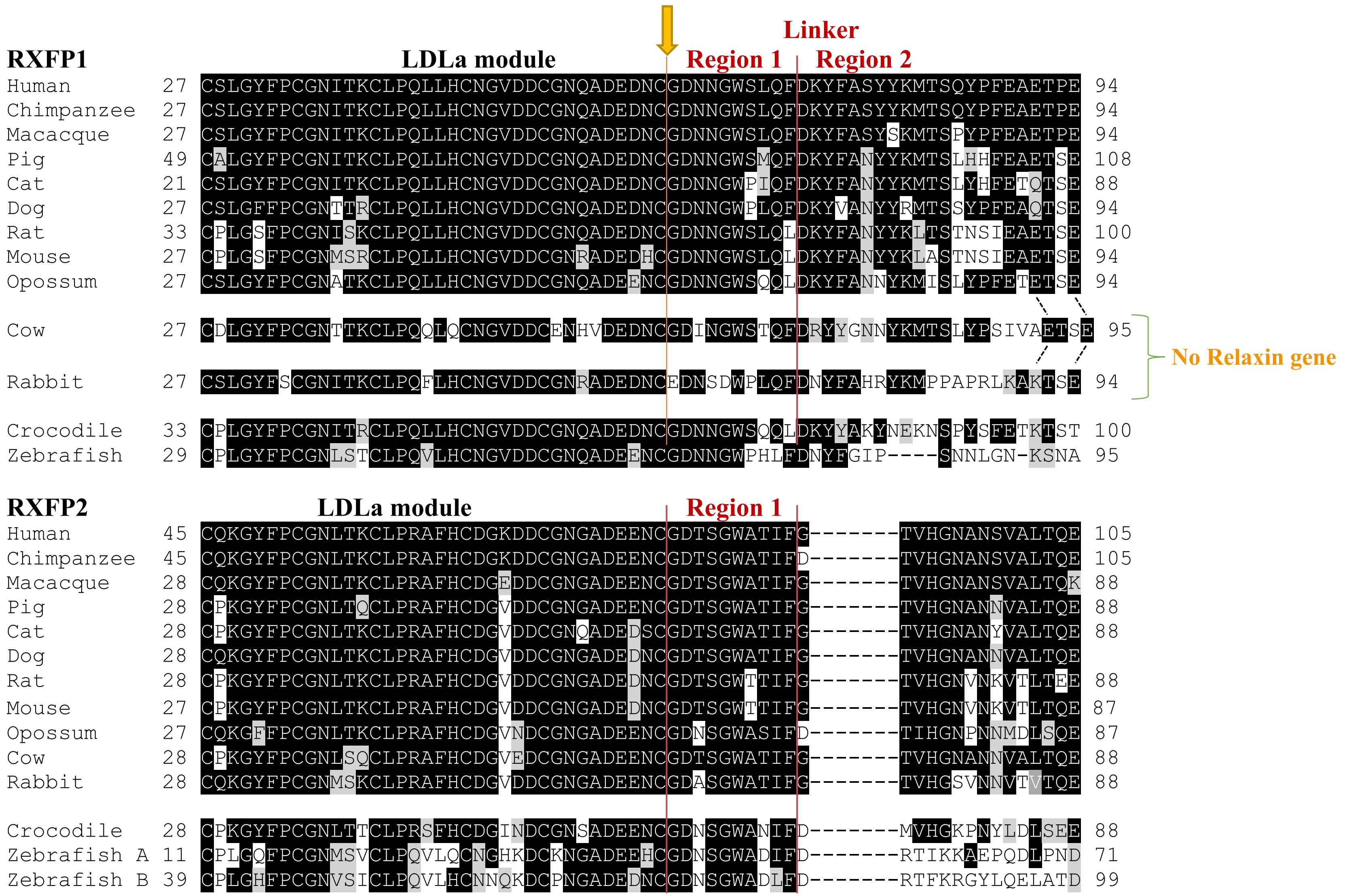

Linker
RXFP1
LDLa module
Region 1
Region 2
Human 27 CSLGYFPCGNITKCLPQLLHCNGVDDCGNQADEDNCGDNNGWSLQFDKYFASYYKMTSQYPFEAETPE 94
Chimpanzee 27 CSLGYFPCGNITKCLPQLLHCNGVDDCGNQADEDNCGDNNGWSLQFDKYFASYYKMTSQYPFEAETPE 94
Macacque 27 CSLGYFPCGNITKCLPQLLHCNGVDDCGNQADEDNCGDNNGWSLQFDKYFASYSKMTSPYPFEAETPE 94
Pig 49 CALGYFPCGNITKCLPQLLHCNGVDDCGNQADEDNCGDNNGWSMQFDKYFANYYKMTSLHHFEAETSE 108
Cat 21 CSLGYFPCGNITKCLPQLLHCNGVDDCGNQADEDNCGDNNGWPIQFDKYFANYYKMTSLYHFETQTSE 88
Dog 27 CSLGFFPCGNTTRCLPQLLHCNGVDDCGNQADEDNCGDNNGWPLQFDKYVANYYRMTSSYPFEAQTSE 94
Rat 33 CPLGSFPCGNISKCLPQLLHCNGVDDCGNQADEDNCGDNNGWSLQLDKYFANYYKLTSTNSIEAETSE 100
Mouse 27 CPLGSFPCGNMSRCLPQLLHCNGVDDCGNRADEDHCGDNNGWSLQLDKYFANYYKLASTNSIEAETSE 94
Opossum 27 CSLGYFPCGNATKCLPQLLHCNGVDDCGNQADEENCGDNNGWSQQLDKYFANNYKMISLYPFETETSE 94
Cow 27 CDLGYFPCGNTTKCLPQQLQCNGVDDCENHVDEDNCGDINGWSTQFDRYYGNNYKMTSLYPSIVAETSE 95
No Relaxin gene
Rabbit 27 CSLGYFSCGNITKCLPQFLHCNGVDDCGNRADEDNCEDNSDWPLQFDNYFAHRYKMPPAPRLKAKTSE 94
Crocodile 33 CPLGYFPCGNITRCLPQLLHCNGVDDCGNQADEDNCGDNNGWSQQLDKYYAKYNEKNSPYSFETKTST 100
Zebrafish 29 CPLGYFPCGNLSTCLPQVLHCNGVDDCGNQADEENCGDNNGWPHLFDNYFGIP----SNNLGN-KSNA 95
RXFP2
LDLa module
Region 1
Human 45 CQKGYFPCGNLTKCLPRAFHCDGKDDCGNGADEENCGDTSGWATIFG--------TVHGNANSVALTQE 105
Chimpanzee 45 CQKGYFPCGNLTKCLPRAFHCDGKDDCGNGADEENCGDTSGWATIFD--------TVHGNANSVALTQE 105
Macacque 28 CQKGYFPCGNLTKCLPRAFHCDGEDDCGNGADEENCGDTSGWATIFG--------TVHGNANSVALTQK 88
Pig 28 CPKGYFPCGNLTQCLPRAFHCDGVDDCGNGADEENCGDTSGWATIFG--------TVHGNANNVALTQE 88
Cat 28 CPKGYFPCGNLTKCLPRAFHCDGVDDCGNQADEDSCGDTSGWATIFG--------TVHGNANYVALTQE 88
Dog 28 CQKGYFPCGNLTKCLPRAFHCDGVDDCGNGADEDNCGDTSGWATIFG--------TVHGNANNVALTQE
Rat 28 CPKGYFPCGNLTKCLPRAFHCDGVDDCGNGADEDNCGDTSGWTTIFG--------TVHGNVNKVTLTEE 88
Mouse 27 CPKGYFPCGNLTKCLPRAFHCDGVDDCGNGADEDNCGDTSGWTTIFG--------TVHGNVNKVTLTQE 87
Opossum 27 CQKGFFPCGNLTKCLPRAFHCDGVNDCGNGADEENCGDNSGWASIFD--------TIHGNPNNMDLSQE 87
Cow 28 CPKGYFPCGNLSQCLPRAFHCDGVEDCGNGADEENCGDTSGWATIFG--------TVHGNANNVALTQE 88
Rabbit 28 CQKGYFPCGNMSKCLPRAFHCDGVDDCGNGADEENCGDASGWATIFG--------TVHGSVNNVTVTQE 88
Crocodile 28 CPKGYFPCGNLTTCLPRSFHCDGINDCGNSADEENCGDNSGWANIFD--------MVHGKPNYLDLSEE 88
Zebrafish A 11 CPLGQFPCGNMSVCLPQVLQCNGHKDCKNGADEEHCGDNSGWADIFD--------RTIKKAEPQDLPND 71
Zebrafish B 39 CPLGHFPCGNVSICLPQVLHCNNQKDCPNGADEDNCGDNSGWADLFD--------RTFKRGYLQELATD 99

**Supplementary Figure 6: Sequence alignment of the LDLa-linker region of RXFP1 and RXFP2 from different species.** Residues GDxxGWxxxF comprise region 1, thought to be important for activation of both RXFP1 and RXFP2 while region 2 is the proposed relaxin-binding site in RXFP1 and is absent from the RXFP2 linker. Species shown are human, chimpanzee (Pan troglodytes), macaque (Macaca Mulatta), pig (Sus scrofa), cat (Felis catus), rat (Rattus norvegicus), Mouse (Mus musculus), opossum (Monodelphis domestica), cow (Bos Taurus), rabbit (Oryctolagus cuniculus), saltwater crocodile (Crocodylus porosus), zebrafish (Danio rerio). Sequences are from Ensembl and UniprotKB. Rat, mouse, macaque and pig RXFP1 have all been shown to respond to relaxin. Cow and rabbit do not have a classic relaxin gene. Crocodile and Zebrafish express a number of relaxin-like peptides with similarities to human relaxin and INSL3 but lack the physiological roles of relaxin and INSL3 in mammals; ie. Mammalian pregnancy and testis descent, respectively. Notably zebrafish RXFP2a and b are expressed in the testis and zebrafish INSL3 is involved in spermatogenesis. The arrow marks the only intron/exon boundary in both the RXFP1 and RXFP2 genes in this sequence region.
